# The N-degron pathway governs autophagy to promote thermotolerance in *Arabidopsis*

**DOI:** 10.1101/2024.07.17.604022

**Authors:** Seu Ha Kim, Jun Seok Park, Myoung-Hoon Lee, Joongyu Seo, Jaekwan Kim, Woo Seok Yang, Jihye Park, Kwangmin Yoo, Jungmin Choi, Jong-Bok Seo, Hyun Kyu Song, Ohkmae K. Park

## Abstract

Autophagy is a vital process that enables plants to adapt to various environmental changes. During heat stress (HS), misfolded and denatured proteins accumulate in cells, necessitating autophagy for their removal. Here, we show that a core autophagy component ATG8a is targeted for degradation via the Arg/N-degron pathway. *ATG8a* is expressed as two alternatively spliced transcripts encoding ATG8a isoforms, namely ATG8a(S) and ATG8a(L), with distinct N-termini. While ATG8a(S) remains stable, ATG8a(L) is N-terminally processed to expose the Arg/N-degron, leading to its degradation. UBR7, identified as an N-recognin, is responsible for ubiquitination and proteasomal degradation of ATG8a(L). Notably, *ATG8a(S)* and *ATG8a(L)* show dynamic expression patterns, fluctuating ATG8a levels during the HS and recovery periods. Our findings highlight the crucial role of ATG8a turnover in conferring thermotolerance, which is governed by Arg/N-degron-mediated regulation. Understanding the molecular basis of ATG8a stability will provide valuable insights into plant resilience to HS under changing climatic conditions.

## INTRODUCTION

N-degron pathways are proteolytic systems that control the half-life of proteins based on the identity of N-terminal residues (Bachmair et al., 1986; Varshavsky, 2011; Gibbs et al., 2014a; Holdsworth et al., 2020). Nascent proteins are synthesized with the N-terminal Met but can be post-translationally processed to expose new N-terminal residues that serve as degradation signals, termed N-degrons, making proteins short-lived *in vivo*. The N-terminal residues are recognized by E3 ubiquitin (Ub) ligase N-recognins that lead to polyubiquitination and proteasomal degradation via the Ub-proteasome system (UPS). In eukaryotes, five N-degron pathways have been defined as: the Arg/N-degron (Varshavsky, 2011; Gibbs et al., 2014a), Ac/N-degron (Hwang et al., 2010; Park et al., 2015), Pro/N-degron (Chen et al., 2017, 2021), formyl-Met/N-degron (Kim et al., 2018), and Gly/N-degron (Timms et al., 2019) pathways. The Arg/N-degron pathway is divided into two branches, in which type I basic (Arg, Lys, His, Cys, Asp, Glu) and type II bulky hydrophobic (Phe, Tyr, Trp, Leu, Ile) N-terminal residues are recognized by the UBR box and ClpS-like domain of N-recognins, respectively (Sriram et al., 2011; Varshavsky, 2019). Arg/N-degrons often arise via enzymatic cascade reactions.

Yeast has a single Arg/N-recognin, Ub ligase N-recognin 1 (UBR1), but the mammalian genome encodes seven UBR box-containing proteins (UBR1-UBR7), of which UBR1, UBR2, UBR4, and UBR5 have been shown to function as N-recognins (Varshavsky, 2011; Tasaki et al., 2012; Gibbs et al., 2014a). In plants, two N-recognins, PROTEOLYSIS 1 (PRT1) and PRT6, have been identified (Potuschak et al., 1998; Garzon et al., 2007). PRT6 contains the UBR box recognizing the type I N-terminal Arg and exhibits high sequence similarity to UBR1 (Kim et al., 2020). PRT1 recognizes type II residues (Phe, Tyr, Trp) via the ZZ domain, which is different from the ClpS-like domain in UBRs. Unlike plants, in yeast and animals, type I and II recognition sites reside within the same UBR protein. N-degron pathways target diverse substrates and regulate a wide array of biological processes (Sriram et al., 2011; Gibbs et al., 2014a; Dissmeyer, 2019; Holdsworth et al., 2020). In plants, substantial progress has been made in the study of the Arg/N-degron pathway, which functions as an O_2_ and NO sensor in response to hypoxia under conditions of waterlogging or submergence (Gibbs et al., 2011; Licausi et al., 2011; Gibbs et al., 2014b; Dissmeyer, 2019).

Macroautophagy (hereafter autophagy) is a highly conserved cellular process in eukaryotes that engulfs cellular components into double membrane-bound autophagosomes and delivers them to the vacuole or lysosome for degradation and recycling (Chung, 2011; Marshall and Vierstra, 2018). Autophagy was initially regarded as a bulk degradation process induced under nutrient deprivation and stress conditions, but it has become clear that it functions to maintain cellular homeostasis by selectively degrading specific targets, such as protein aggregates and damaged organelles, in a process termed selective autophagy (Zaffagnini and Martens, 2016). In animal systems, the autophagic adaptor p62 acts as an N-recognin mediating crosstalk between UPS and autophagy (Cha-Molstad et al., 2015, 2017; Yoo et al., 2018; Heo et al., 2021). Autophagosome formation is an essential part of autophagy and requires core autophagy machinery composed of autophagy-related (ATG) proteins. Molecular and genetic analyses of autophagy-defective *atg* mutants indicate that autophagy plays important roles in development and abiotic and biotic stress responses (Wang et al., 2018).

Most ATG proteins assemble into complexes and are recruited to the phagophore assembly site (PAS) to mediate a series of autophagic events (Ichimura et al., 2000; Yoshimoto et al., 2004; Feng et al, 2014; Marshall and Vierstra, 2018; Martens and Fracchiolla, 2020). In addition to its role in phagophore expansion, ATG8 is essential for cargo selection in selective autophagy through its interaction with cargo receptors such as Neighbor of Brca1 (NBR1) (Bu et al., 2020). In contrast to a single ATG8 in yeast, ATG8 is diversified in higher eukaryotes. For example, *Arabidopsis* has nine ATG8 isoforms (ATG8a-i). While the reason of ATG8 diversity remains elusive, ATG8 variants may have specificity for cargo receptors to target distinct cargos in selective autophagy (Kellner et al., 2017). Despite substantial studies into autophagosome biogenesis and its components, the dynamic features of autophagy are not fully understood. Here we report that the Arg/N-degron pathway regulates temporal ATG8a turnover during heat stress (HS) responses, which is a crucial process ensuring plant survival in fluctuating thermal environments.

## RESULTS

### ATG8a is regulated by the Arg/N-degron pathway

To explore the role of ATG8, we expressed two *Arabidopsis* ATG8 isoforms, ATG8a and ATG8e, fused with either an N-terminal MYC tag (MYC-ATG8a/8e) or a C-terminal hemagglutinin (HA) tag (ATG8a/8e-HA) in wild-type Col-0 protopalsts. Unlike ATG8e, ATG8a showed unexpectedly low expression, irrespective of the type and position of affinity tags (Figure 1A). This intriguing observation led us to suspect that ATG8a might be degraded by the UPS. To test this, we monitored ATG8a expression in the presence of the proteasome inhibitor MG132. Immunoblotting with antibodies against the HA tag and ATG8a revealed that MG132 treatment increases ATG8a protein levels. However, MYC-ATG8a proteins remained barely detectable by the anti-MYC antibody (Figure 1B), suggesting that the N-terminal part of ATG8a may be lost. Untagged ATG8a proteins were also stabilized by MG132 treatment (Figure 1C). Therefore, affinity tags are unlikely to interfere with ATG8a expression. The specificity of the anti-ATG8a antibody for ATG8a was validated by immunoblot analysis using purified recombinant proteins of ATG8 isoforms (ATG8a-i) (Figure S1A), excluding the possibility of cross-reactivity with other ATG8 isoforms in *Arabidopsis*. These immunoblotting data suggest that ATG8a undergoes 26S proteasome-dependent degradation, potentially involving the truncation of its N-terminal part. The degradation patterns of ATG8a were similar in protoplasts from wild-type plants grown under both long-day and short-day conditions, indicating that ATG8a stability is not influenced by photoperiods (Figure S1B).

**Figure 1.**
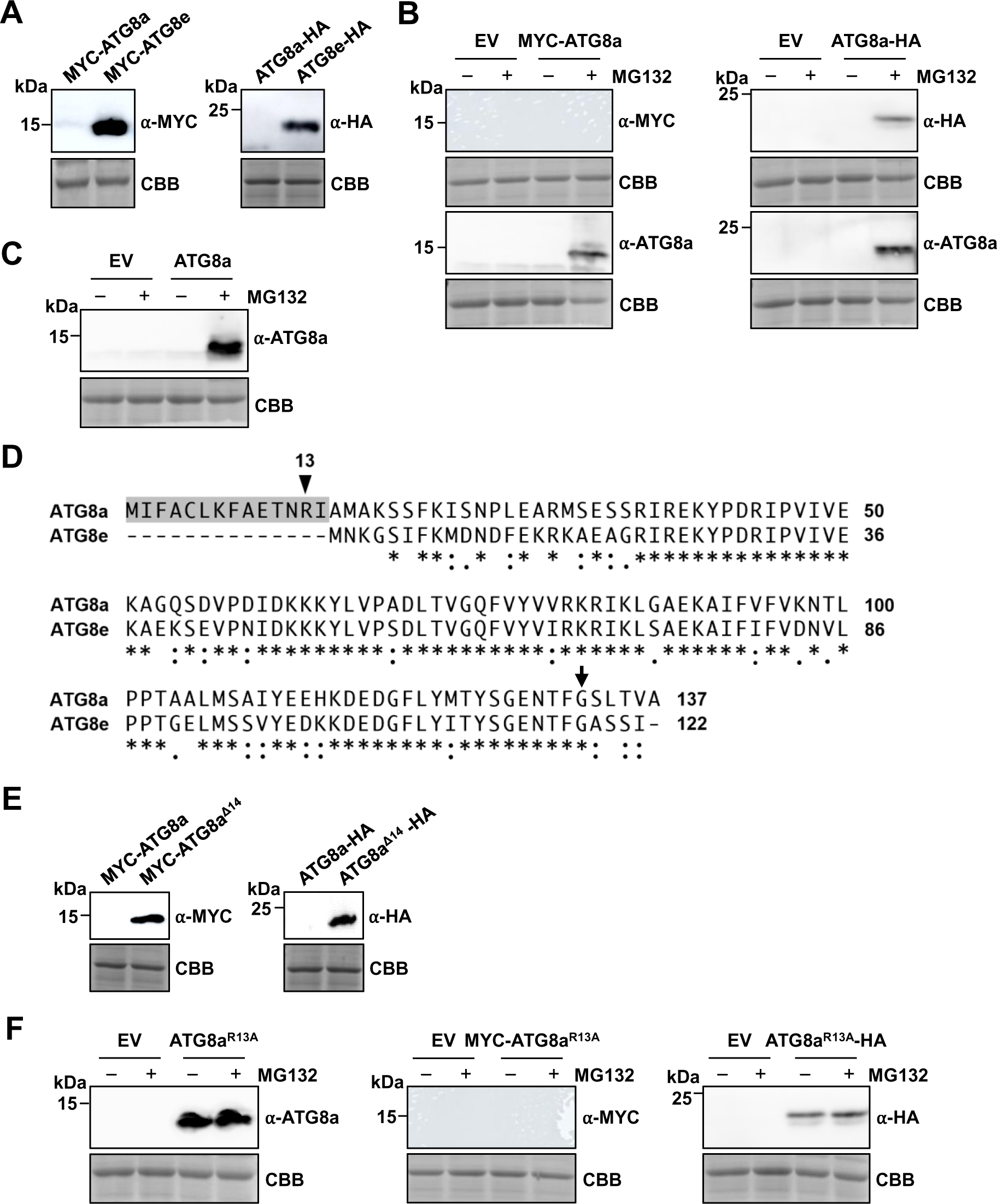
ATG8a undergoes Arg/N-degron-mediated degradation. (A) ATG8a expression is barely detectable in Col-0 protoplasts. MYC-ATG8a/ATG8e and ATG8a/ATG8e-HA were expressed in Col-0 protoplasts. (B and C) ATG8a either with (B) or without (C) affinity tags undergo 26S proteasome-dependent degradation. MYC-ATG8a, ATG8a-HA, and untagged ATG8a were expressed in Col-0 protoplasts, followed by treatments with cycloheximide (100 μM) and MG132 (10 μM) for 3 h. (D) Sequence alignment of ATG8a and ATG8e shows that ATG8a has 14 additional amino acids at the N-terminus compared to ATG8e. Sequences were aligned using Clustal Omega online (https://www.ebi.ac.uk/Tools/msa/clustalo/). The N-terminal 14 residues are shaded, and the putative N-degron residue R^13^ is indicated by an arrowhead in ATG8a. The Gly residue exposed at the C-terminus after C-terminal cleavage and conjugated to PE is indicated by an arrow. Consensus symbols are as follows: asterisks indicate identical residues; colons and periods indicate conserved and semiconserved substitutions, respectively. (E) Deletion of N-terminal 14 residues stabilizes ATG8a. MYC-ATG8a/ATG8a^Δ14^ and ATG8a/ATG8a^Δ14^-HA were expressed in Col-0 protoplasts. (F) R^13^A mutation stabilizes ATG8a. ATG8a^R13A^ without affinity tags, MYC-ATG8a^R13A^, and ATG8a^R13A^-HA were expressed in Col-0 protoplasts, followed by treatments with cycloheximide (100 μM) and MG132 (10 μM) for 3 h. ATG8a proteins were analyzed by immunoblotting with respective antibodies, and Coomassie brilliant blue (CBB) staining served as a loading control. EV, empty vector.

Alignment of ATG8a and ATG8e sequences revealed that ATG8a has 14 additional amino acids at the N-terminus compared to ATG8e, suggesting that these extra residues might contribute to the stability difference between ATG8a and ATG8e (Figure 1D). To investigate this, we expressed an ATG8a variant with the N-terminal 14 residues deleted (ATG8a^Δ14^) and found that this deletion stabilizes ATG8a (Figure 1E). This N-terminus-dependent ATG8a degradation led to the plausible assumption that ATG8a contains an N-degron and is regulated by the N-degron pathway. Notably, the N-terminal sequence of ATG8a features characteristics of an Arg/N-degron, with Arg at position 13 followed by a hydrophobic residue Ile at position 14 (Figure 1D) (Choi et al., 2010). Indeed, Arg 13 to Ala mutation stabilized ATG8a^R13A^, while MYC-ATG8a^R13A^ remained barely detectable by the anti-MYC antibody (Figure 1F). These results suggest that ATG8a exposes Arg at the N-terminus as a primary destabilizing residue after cleavage before R^13^ and undergoes Arg/N-degron-mediated degradation.

ATG8 undergoes phosphatidylethanolamine (PE)-conjugation at a Gly (G^132^ in ATG8a; Figure 1D) exposed at the C-terminus after C-terminal cleavage during autophagy activation (Woo et al., 2014). To investigate whether ATG8a lipidation affects its expression and stability, we generated a non-lipidated (NL) ATG8a mutant, ATG8aNL, with FGS^131-133^ to AAA changes. We then examined the expression of both tagged and untagged ATG8aNL proteins (Figure S1C). No differences in expression patterns were observed between wild-type and NL forms of ATG8a. This implies that a significant portion of ATG8a exists as a free cytosolic form under normal conditions.

### ATG8a is N-terminally cleaved to expose the Arg/N-degron

To investigate the N-terminal truncation of ATG8a, residues around R^13^ of MYC-ATG8a were mutated to determine whether these mutant proteins are detectable with an anti-MYC antibody (Figure 2A). Unlike ATG8a^AVG^ with RIA^13-15^ to AVG substitutions, ATG8a^7A^ with mutations in 7 residues (positions 10-16) around R^13^, and ATG8a^3A^ with mutations in 3 residues (positions 10-12) before R^13^ were stably detected by the anti-MYC antibody (Figure 2B). Additional mutational analysis of these three residues showed that two residues, T^11^ and N^12^, preceding R^13^ are critical for N-terminal cleavage of ATG8a (Figure 2C). These results are consistent with earlier studies showing that the residues N-terminal to the cleavage site in substrate proteins largely determine the specificity of proteolytic enzymes (Qi et al., 2017).

**Figure 2.**
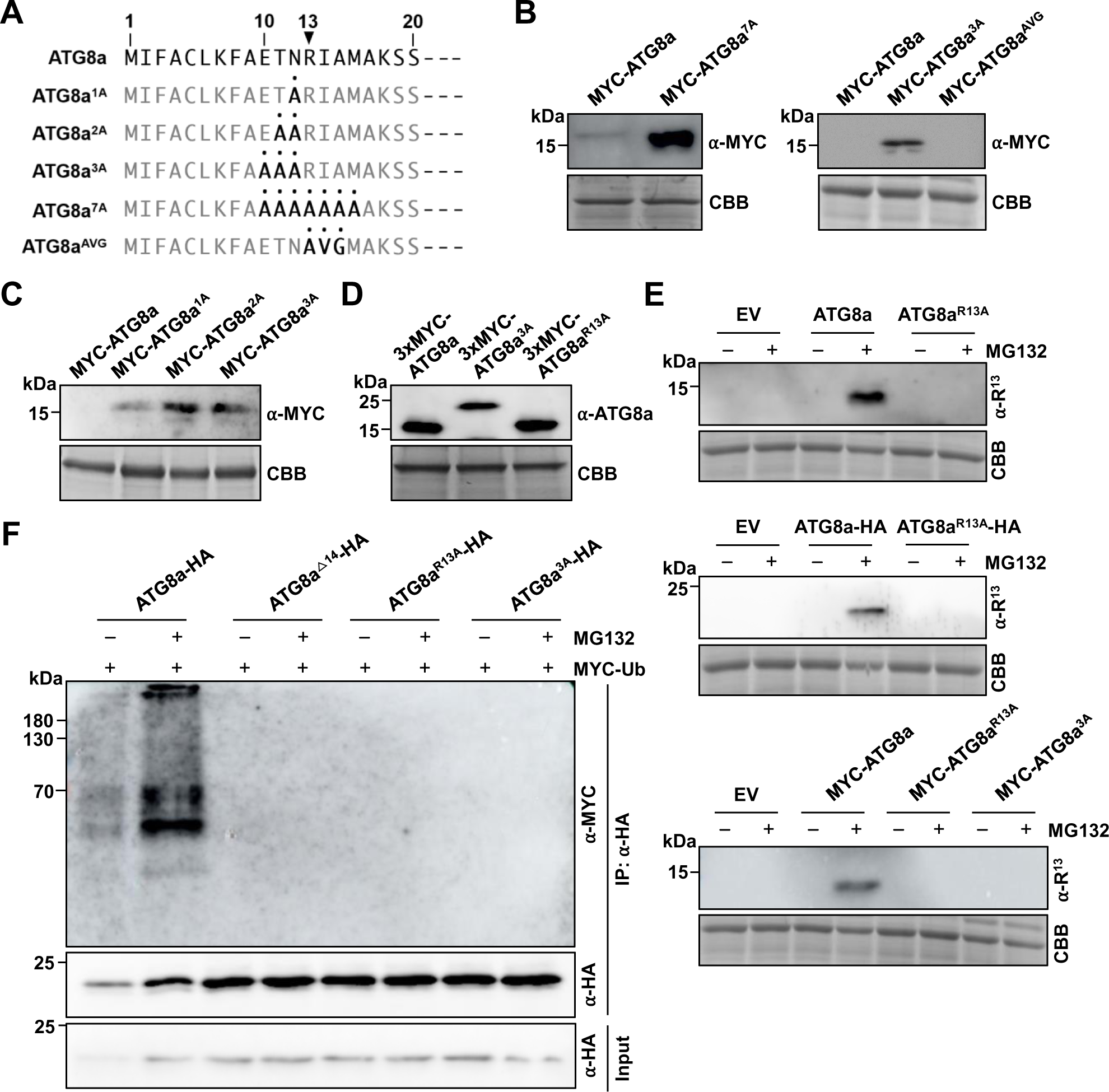
ATG8a is N-terminally cleaved to expose the Arg/N-degron. (A) N-terminal sequences of ATG8a mutants with the indicated substitutions. (B) Mutations at residues preceding R^13^ stabilize ATG8a. MYC-tagged ATG8a, ATG8a^7A^, ATG8a^3A^, and ATG8a^AVG^ were expressed in Col-0 protoplasts. (C) Mutations at T^11^ and N^12^ stabilize ATG8a. MYC-tagged ATG8a, ATG8a^1A^, ATG8a^2A^, and ATG8a^3A^ were expressed in Col-0 protoplasts. (D) ATG8a is N-terminally processed. ATG8a, ATG8a^3A^, and ATG8a^R13A^ fused with 3 MYC tags (3xMYC) at the N-terminus were expressed in Col-0 protoplasts, followed by treatment with MG132 (10 μM) for 3 h. (E) ATG8a exposes R^13^ at the N-terminus. ATG8a, ATG8a^R13A^, and ATG8a^3A^, either with or without affinity tags, were expressed in Col-0 protoplasts, followed by treatments with cycloheximide (100 μM) and MG132 (10 μM) for 3 h. EV, empty vector; α-R^13^, anti-R^13^- ATG8a antibody. (F) *In vivo* ubiquitination assay for ATG8a polyubiquitination. ATG8a, ATG8a^Δ14^, ATG8a^R13A^, and ATG8a^3A^ fused with HA at the C-terminus were expressed together with MYC-Ub in Col-0 protoplasts, followed by treatment with MG132 (10 μM) for 3 h. Protein lysates were subjected to immunoprecipitation with the anti-HA antibody. Input shows 2% of the amount used in reactions. IP, immunoprecipitation. ATG8a proteins were analyzed by immunoblotting with respective antibodies, and Coomassie brilliant blue (CBB) staining served as a loading control.

When ATG8a fused with three MYC tags (3xMYC) at the N-terminus was expressed in the presence of MG132 and immunoblotted with the anti-ATG8a antibody, it clearly revealed a decrease in the molecular size of ATG8a and ATG8^R13A^ compared to ATG8a^3A^ by approximately 5 kDa, corresponding to 3xMYC and N-terminal 12 residues (Figure 2D), supporting the N-terminal processing of ATG8a. To verify that ATG8a exposes R^13^ at the N-terminus, we raised the anti-R^13^-ATG8a antibody that specifically recognizes R^13^-ATG8a using the peptide RIAMAKSSFKI^13-23^. We also prepared R^13^-ATG8a and A^13^-ATG8a with an Arg 13 to Ala substitution using the LC3B-fusion technique (Kim et al., 2020). Immunoblot analysis using R^13^-ATG8a and A^13^-ATG8a and the peptide competition assay confirmed the antibody specificity for R^13^-ATG8a (Figure S1D). As predicted, the anti-R^13^-ATG8a antibody immunoblotted both untagged and tagged ATG8a stabilized by MG132 treatment, but not ATG8a^R13A^ and ATG8a^3A^ (Figure 2E).

We also tested for ATG8a ubiquitination in an *in vivo* ubiquitination assay in protoplasts by co-expressing MYC-tagged Ub. Consistently, high-molecular weight smears, indicative of polyubiquitination, were observed for ATG8a, but not for ATG8a^Δ14^, ATG8^R13A^, and ATG8a^3A^ which were resistant to proteasomal degradation (Figure 2F). Collectively, our results suggest that N-terminally truncated R^13^-ATG8a is a target substrate of the Arg/N-degron pathway.

### UBR7 functions as an N-recognin for ATG8a

Next, we sought to identify an N-recognin that targets ATG8a to proteasomal degradation via the Arg/N-degron pathway. *Arabidopsis* has three UBR box-containing proteins, PRT6, BIG (Gil et al., 2001), and AT4G23860 (Holdsworth et al., 2020). AT4G23860, which shows considerable similarity to mammalian UBR7, is hereafter referred to as *Arabidopsis* UBR7 (Figure S2A). We examined whether ATG8a stability is regulated by PRT6, BIG, or UBR7. In transient expression assays, ATG8a-HA did not accumulate without MG132 treatment in protoplasts of *prt6-1* and *big-3* knockout mutants, similar to wild-type protoplasts (Figure 3A). Since PRT6 is a known N-recognin, an artificial R-GUS substrate with the N-terminal Arg was generated through the Ub fusion technique (Bachmair et al., 1986), and its degradation was tested in *prt6-1* protoplasts, as previously reported (Garzon et al., 2007). R-GUS proteins, but not M-GUS, underwent proteasomal degradation in wild-type protoplasts, while both R- and M-GUS accumulated in *prt6-1* (Figure S2B). These results imply that the N-terminal Arg of N-degron substrates is required but not sufficient for specific binding to N-recognins. Notably, ATG8a-HA levels were elevated in *ubr7* protoplasts regardless of MG132 treatment (Figures 3B and S2C-E). Similarly, R^13^-ATG8a, generated through the Ub fusion technique, was degraded in wild-type protoplasts but stabilized in *ubr7*, unlike A^13^-ATG8a, which was stably expressed (Figure 3C). These results suggest that UBR7 is the cognate N-recognin for R^13^- ATG8a.

**Figure 3.**
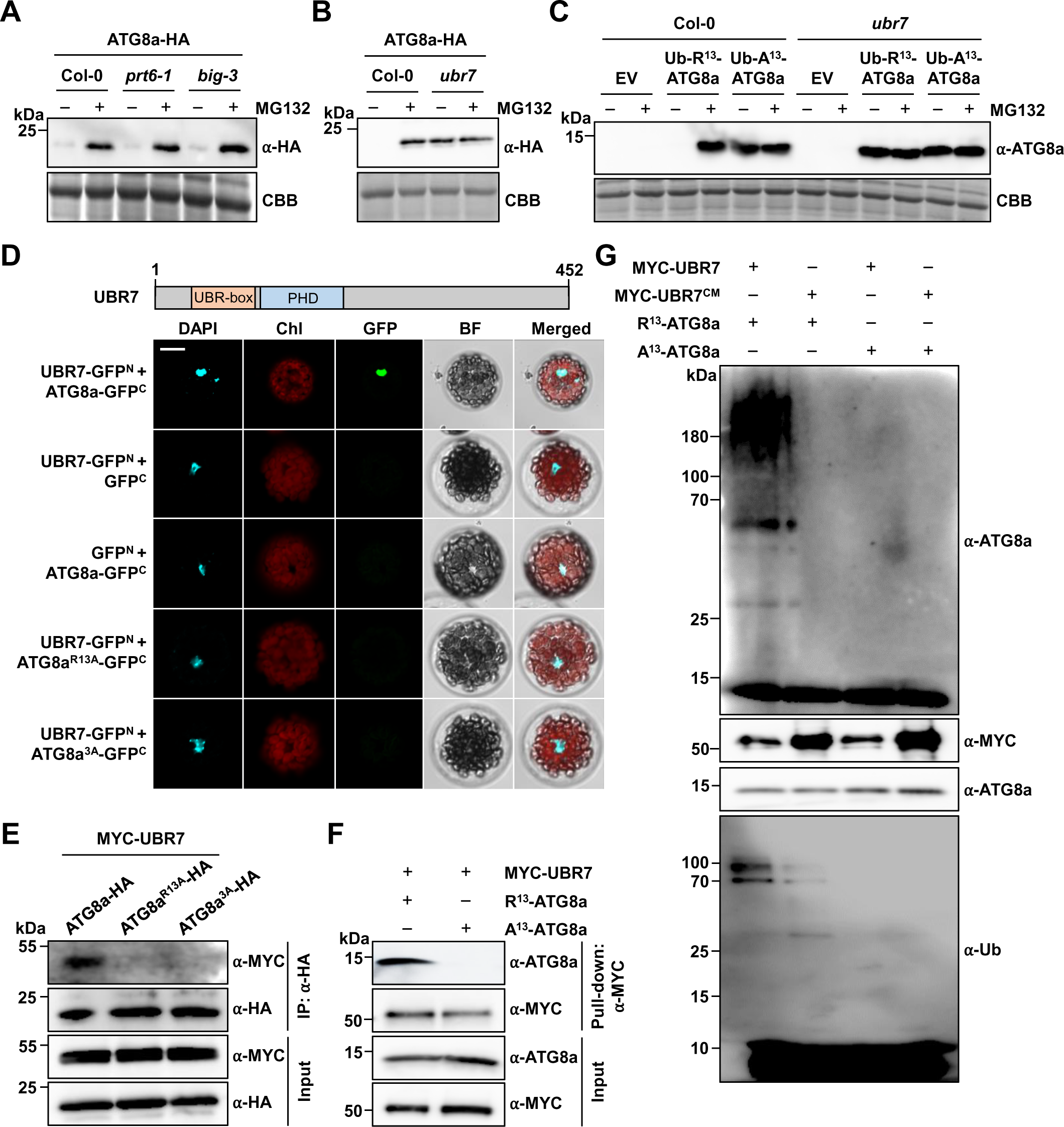
UBR7 is an E3 Ub ligase N-recognin for ATG8a. (A and B) ATG8a expression is not altered in *prt6-1* and *big-3* (A) but is elevated in *ubr7* (B) mutants. ATG8a-HA was expressed in Col-0, *prt6-1*, *big-3*, and *ubr7* protoplasts, followed by treatments with cycloheximide (100 μM) and MG132 (10 μM) for 3 h. (C) R^13^-ATG8a generated through the Ub fusion technique is degraded in wild-type but stabilized in *ubr7* mutant. Ub-R^13^/A^13^-ATG8a were expressed in Col-0 and *ubr7* protoplasts, followed by treatments with cycloheximide (100 μM) and MG132 (10 μM) for 3 h. EV, empty vector. (D) BiFC assay for *in vivo* interaction between ATG8a and UBR7. Domain structure of *Arabidopsis* UBR7 is shown (top). GFP^N^, GFP^C^, and their fusions with UBR7, ATG8a, ATG8a^R13A^, and ATG8a^3A^ were co-expressed in Col-0 protoplasts, followed by treatment with MG132 (10 μM) for 3 h. Reconstituted GFP fluorescence was visualized under a confocal microscope. DAPI staining indicates the location of nuclei. Chl, chlorophyll; BF, bright field. Bars, 10 μm. (E) *In vivo* co-immunoprecipitation assay for interaction between ATG8a and UBR7. Protein lysates were prepared from Col-0 protoplasts co-expressing MYC-UBR7 and ATG8a-HA, ATG8a^R13A^, or ATG8a^3A^ and subjected to immunoprecipitation with the anti-HA antibody. Input shows 2% of the amount used in binding reactions. IP, immunoprecipitation. (F) *In vitro* pull-down assay for interaction between ATG8a and UBR7. MYC-UBR7 was incubated with R^13^- ATG8a or A^13^-ATG8a and pulled down by incubation with the anti-MYC antibody and protein G agarose beads. Input shows 5% of the amount used in binding reactions. (G) *In vitro* ubiquitination assay for UBR7-mediated R^13^-ATG8a polyubiquitination. MYC-UBR or MYC-UBR7^CM^ was incubated with either R^13^-ATG8a or A^13^-ATG8a in the presence of Ub, human E1, and *Arabidopsis* E2 enzymes. ATG8a and UBR7 proteins were analyzed by immunoblotting with respective antibodies, and Coomassie brilliant blue (CBB) staining served as a loading control.

To assess the interaction between ATG8a and UBR7, bimolecular fluorescence complementation (BiFC) assays were conducted by co-expressing UBR7 or the UBR box domain of UBR7 fused with the N-terminal half of GFP (GFP^N^) and ATG8a, ATG8a^R13A^, and ATG8a^3A^ fused with the C-terminal half of GFP (GFP^C^) under the CaMV 35S promoter in protoplasts (Figure S2F). GFP signals were detected only in protoplasts co-expressing UBR7/UBR box and ATG8a, while both UBR7 and UBR box domain showed no fluorescent signals with ATG8a^R13A^ and ATG8a^3A^ (Figures 3D and S2G). It was noted that UBR7 is expressed and interacts with ATG8a only in the nucleus, as overlapped with the nuclear 4′,6-diamidino-2-phenylindole (DAPI) signal, although both ATG8a and ATG8aNL are localized in both the cytoplasm and nucleus (Figures 3D and S3). Their interaction was further evaluated by a co-immunoprecipitation assay in protoplasts. Consistently, UBR7 co-immunoprecipitated with ATG8a, but not with ATG8a^R13A^ and ATG8a^3A^ (Figure 3E). Moreover, it was confirmed that UBR7 binds specifically to the N-terminal Arg of ATG8a using R^13^-ATG8a and A^13^-ATG8a proteins prepared by the LC3B-fusion technique (Kim et al., 2020). An *in vitro* pull-down assay clearly showed a specific interaction between MYC-UBR7 and R^13^-ATG8a (Figure 3F). Since mammalian UBR7 was previously shown to have E3 Ub ligase activity (Adhikary et al., 2019), we investigated whether *Arabidopsis* UBR7 functions as an E3 Ub ligase N-recognin. MYC-UBR7 and its catalytic mutant MYC-UBR7^CM^ with H^157^S and H^160^S substitutions (Adhikary et al., 2019) were incubated with R^13^-ATG8a and A^13^-ATG8a in an *in vitro* ubiquitination assay. The results indicated that UBR7 can polyubiquitinate R^13^-ATG8a but not A^13^-ATG8a (Figure 3G). E3 Ub ligase activity was not detected for UBR7^CM^. Together, these results demonstrate that UBR7, acting as an N-recognin, recognizes and polyubiquitinates ATG8a with Arg exposed at its N-terminus.

### ATG8a is expressed as splice variants *ATG8*(*S*) and *ATG8a*(*L*)

While investigating the biological significance of ATG8a regulation by the Arg/N-degron pathway, we had an intriguing finding in the genomic structure that *ATG8a* undergoes alternative splicing (AS) and is expressed as two splice variants, AT4G21980.1 and AT4G21980.2 (Figure 4A). AT4G21980.2 is an intron retention (IR) variant that encodes ATG8a with an additional 15 amino acids at the N-terminus, bearing the N-degron; AT4G21980.1 and AT4G21980.2 are hereafter designated as *ATG8*(*S*) and *ATG8a*(*L*), respectively. Since ATG8a(L) expression leads to the elimination of ATG8a proteins, differential generation of these splice variants may be a mechanism to modulate ATG8a protein levels.

**Figure 4.**
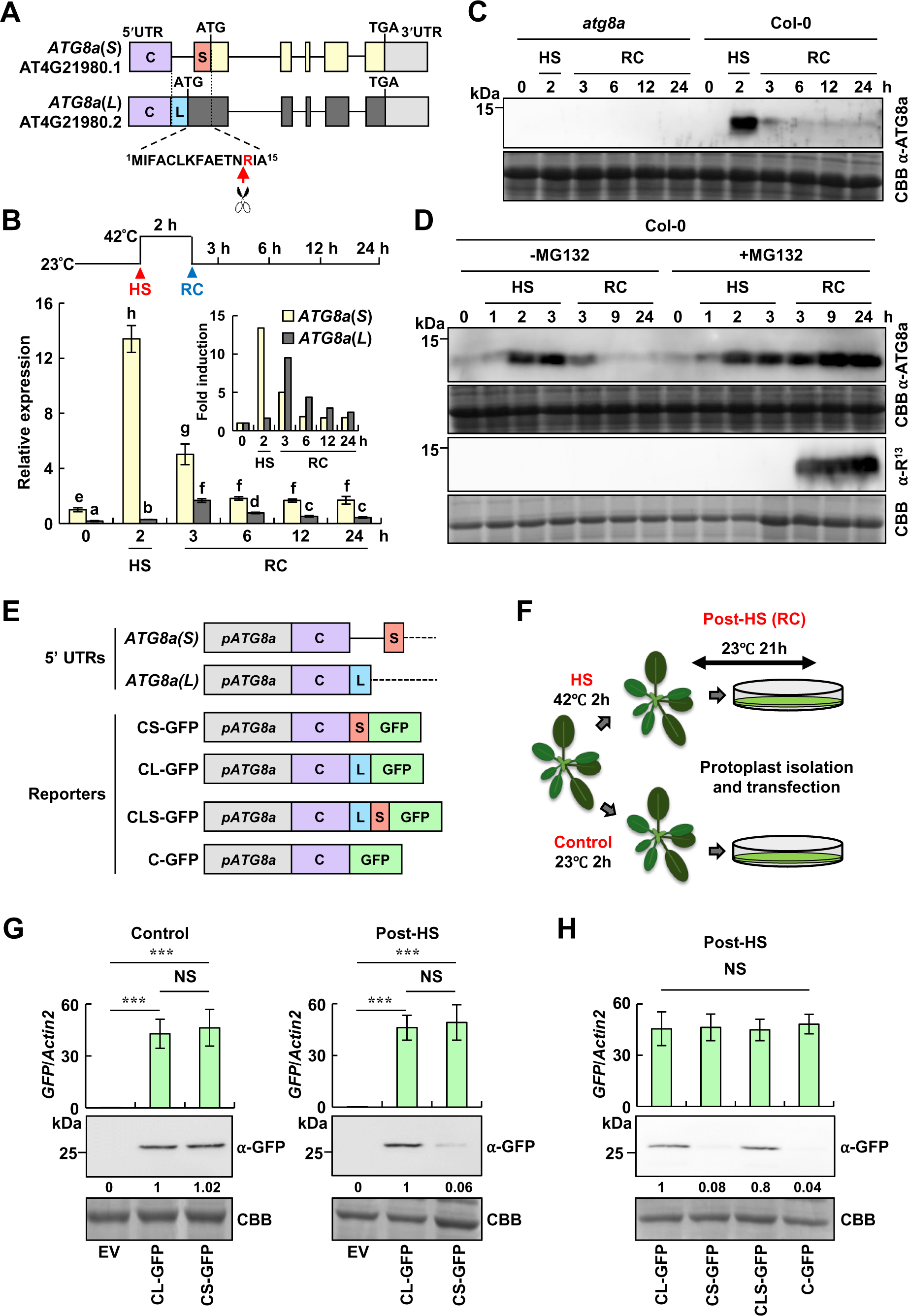
R^13^-ATG8a is expressed and degraded during HS recovery. (A) Schematic diagram of ATG8a splice variants, *ATG8*(*S*) and *ATG8a*(*L*). Exons are indicated by yellow and dark gray (coding) and purple, red, blue, and light gray (UTRs) boxes. Introns are shown in lines. The additional N-terminal 15 residues in ATG8a(L) are shown at the bottom. (B) *ATG8*(*S*) and *ATG8a*(*L*) are differentially expressed during HS responses. *ATG8*(*S*) and *ATG8a*(*L*) expression were analyzed by RT-qPCR in Col-0 plants exposed to HS (42°C) for 2 h and recovery (23°C) for the indicated times. Data represent means ± SD (*n* = 4 biological replicates). Different letters indicate significant differences (Tukey’s HSD test; *P* < 0.05). (C) ATG8a abundance changes during HS responses. ATG8a expression was monitored during HS and recovery in Col-0 and *atg8a* plants. (D) ATG8a undergoes proteasomal degradation during HS recovery. ATG8a expression was monitored in the presence or absence of MG132 during HS and recovery in Col-0 plants. For MG132 treatment, plants were sprayed with MG132 (10 μM) prior to exposure to HS and recovery conditions. α-R^13^, anti-R^13^-ATG8a antibody. (E) Schematic diagram of reporter constructs used for translational activity. 5’ UTRs of *ATG8*(*S*) and *ATG8a*(*L*) are shown, containing distinct sequences indicated as S and L, respectively, in addition to the common sequence C (top). The reporter constructs carry the native promoter *pATG8a* and different combinations of C, S, and L, fused to the *GFP* gene. (F) Scheme of HS treatment and post-HS (recovery) in protoplasts. Col-0 plants were exposed to HS at 42°C for 2 h and subjected to reporter assays in protoplasts. Protoplast isolation and transfection were performed at 23°C, and therefore this period was regarded as the post-HS. (G and H) *ATG8a*(*L*)-specific L sequence stimulates translation under post-HS. Control (no HS treatment) and post-HS protoplasts were transfected with reporter constructs carrying *ATG8a*(*L*) 5’ UTR (CL-GFP) and *ATG8a*(*S*) 5’ UTR (CS-GFP) (G) and different combinations of C, S, and L in 5’ UTRs (H). *GFP* transcript (top) and GFP protein (bottom) levels were determined by RT-qPCR and immunoblotting with the anti-GFP antibody, respectively. In RT-qPCR (top), *Actin2* was used as a control. Data represent means ± SD (*n* = 4 biological replicates). Asterisks indicate significant differences between EV and reporter constructs (*t* test; \*\*\**P* < 0.001). In immunoblotting, relative intensities of GFP normalized to the loading control are shown as numerical values. NS, not significant; EV, empty vector. ATG8a proteins were analyzed by immunoblotting with respective antibodies, and Coomassie brilliant blue (CBB) staining served as a loading control. HS, heat stress; RC, recovery.

### ATG8a(L) expression leads to N-degron-mediated ATG8a degradation during thermorecovery

Considering that autophagy is critical for plant adaptation to environment, we examined whether Arg/N-degron-mediated ATG8a(L) regulation contributes to stress responses. For this, we analyzed the expression patterns of ATG8(S) and ATG8a(L) in *Arabidopsis* Col-0 plants during stress responses with three criteria: (1) induction of *ATG8a*(*L*) expression, (2) concurrent decrease in ATG8a protein levels, and (3) restoration of reduced ATG8a levels with MG132 treatment. Given that autophagy is important for clearing protein aggregates, we investigated its relevance to endoplasmic reticulum (ER) stress caused by the accumulation of unfolded and misfolded proteins. The expression of *ATG8*(*S*) and *ATG8a*(*L*) was monitored in plants after treatments with dithiothreitol (DTT) and tunicamycin (TM), two common ER stress inducers (Back et al., 2005). The transcript level of *ATG8a*(*L*) was much lower than that of *ATG8*(*S*), and DTT and TM treatments gradually increased the expression of *ATG8*(*S*) over 48 and 72 h, respectively (Figures S4A and S4B). In contrast, ATG8a protein levels decreased after peaking at 12 and 48 h, respectively, which was not significantly affected by MG132 treatment (Figures S4C and S4D). These results suggest that ATG8a(L) degradation via the N-degron pathway is not implicated in ER stress responses.

HS is a detrimental abiotic stress that also leads to the accumulation of heat-denatured protein aggregates. Since autophagy is important for thermotolerance and HS recovery (Zhou et al., 2013; Sedaghatmehr et al., 2019), we analyzed the expression of *ATG8*(*S*) and *ATG8a*(*L*) during HS and recovery (Figure 4B). *ATG8*(*S*) was highly expressed in plants exposed to high temperature (42°C) but decreased during HS recovery. In contrast, *ATG8a*(*L*) showed rapid induction after release from HS (Figure 4B, inset), but its abundance was still three-fold less than that of *ATG8*(*S*). We then determined protein levels of ATG8a. ATG8a proteins accumulated under HS but largely decreased after HS (Figure 4C). ATG8a proteins were barely detected in *atg8a* mutant (Figure 4C), further verifying the specificity of the anti-ATG8a antibody. Notably, MG132 treatment restored the abundance of ATG8a during HS recovery, which corresponded to R^13^-ATG8a, as revealed by immunoblotting with the anti-R^13^-ATG8a antibody (Figure 4D). These expression data suggest that ATG8a(S) and ATG8a(L) are major forms expressed during HS and recovery, respectively, and that ATG8a(L) undergoes Arg/N-degron-mediated degradation and is responsible for a large reduction of ATG8a during the post-HS period.

### *ATG8a*(*L*) 5’ UTR promotes translation of *ATG8a*(*L*) during thermorecovery

It was puzzling because *ATG8*(*S*) transcripts accumulated at a much higher level than *ATG8a*(*L*), making it difficult for ATG8a(L) to be expressed as the major form during HS recovery. Therefore, we reasoned that the production of ATG8a(S) and ATG8a(L) may be regulated by an additional post-transcriptional mechanism. Of note, *ATG8*(*S*) and *ATG8a*(*L*) have distinct 5’ untranslated regions (UTRs), in which the 5’ part is common (designated as C) and the 3’ part contains sequences specific to each (designated as S and L, respectively) (Figures 4A and 4E).

To examine the translational activity of 5’ UTRs of *ATG8*(*S*) and *ATG8a*(*L*), we constructed native promoter (*pATG8a*)-driven green fluorescent protein (GFP) reporters, *pATG8a*:CS-GFP and *pATG8a*:CL-GFP, with *GFP* preceded by 5’ UTRs of *ATG8*(*S*) and *ATG8a*(*L*), respectively (Figure 4E). GFP reporter assays were performed by introducing the constructs into protoplasts from wild-type plants that were either unexposed (control) or exposed to HS for 2h (Figure 4F). Since protoplast isolation and transfection proceeded at 23°C, this period was regarded as the post-HS recovery phase. As determined by immunoblotting with the anti-GFP antibody, the translational activity of *ATG8a*(*L*) 5’ UTR (CL) was approximately 17-fold higher than that of *ATG8a*(*S*) 5’ UTR (CS) in post-HS protoplasts, while this difference was not shown in control protoplasts (Figure 4G).

To further evaluate whether *ATG8*(*S*)- and *ATG8a*(*L*)-specific S and L sequences in 5’ UTRs are critical for differential translation of ATG8(S) and ATG8a(L) during HS recovery, we constructed *pATG8a*:CLS-GFP with both L and S sequences and *pATG8a*:C-GFP with only the common C sequence (Figure 4E). Their expression was then compared with that of *pATG8a*:CS-GFP and *pATG8a*:CL-GFP (Figure 4H). In post-HS protoplasts, CLS-GFP translation was stimulated to a level close to that of CL-GFP, while C-GFP was expressed at a much lower level, comparable to that of CS-GFP (Figure 4H). *GFP* transcripts accumulated at similar levels across different constructs (Figures 4G and 4H). These results suggest that the L sequence of *ATG8a*(*L*) 5’ UTR has a stimulatory effect on translation of *ATG8a*(*L*) and significantly increases ATG8a(L) abundance during HS recovery.

### ATG8a degradation is UBR7-dependent during thermorecovery

To further validate whether ATG8a(L) is responsible for ATG8a degradation during the HS recovery period, we generated *ATG8a*(*S*)-expressing *pATG8:ATG8a*(*S*)/*atg8a* complementation lines, in which the native promoter-driven *ATG8a*(*S*) is introduced into *atg8a* mutant, and subjected them to HS and recovery conditions (Figure 5A). The ATG8a expression remained high during both the HS and recovery phases in these complementation lines, implying that the decrease in ATG8a abundance after HS is due to ATG8a(L) production, followed by subsequent degradation. To determine whether ATG8a degradation is UBR7-dependent, further tests were conducted in *ubr7* mutant and *UBR7*-silenced lines (TRV2-UBR7) generated by tobacco rattle virus (TRV)-based virus-induced gene silencing (VIGS) (Burch-Smith et al., 2006; Yang et al., 2021). VIGS was successful, leading to a 56-96% reduction of *UBR7* transcripts in TRV2-UBR7 lines compared to the TRV2 control, with significant induction of *UBR7* during thermorecovery (Figure 5B). Consistently, no reduction in ATG8a protein levels was observed in both *ubr7* and TRV2-UBR7 plants, and R^13^-ATG8a proteins accumulated after release from HS (Figures 5C and 5D). These results demonstrate that the decrease in ATG8a protein abundance after HS is due to ATG8a(L) expression, which occurs through the UBR7-mediated N-degron pathway.

**Figure 5.**
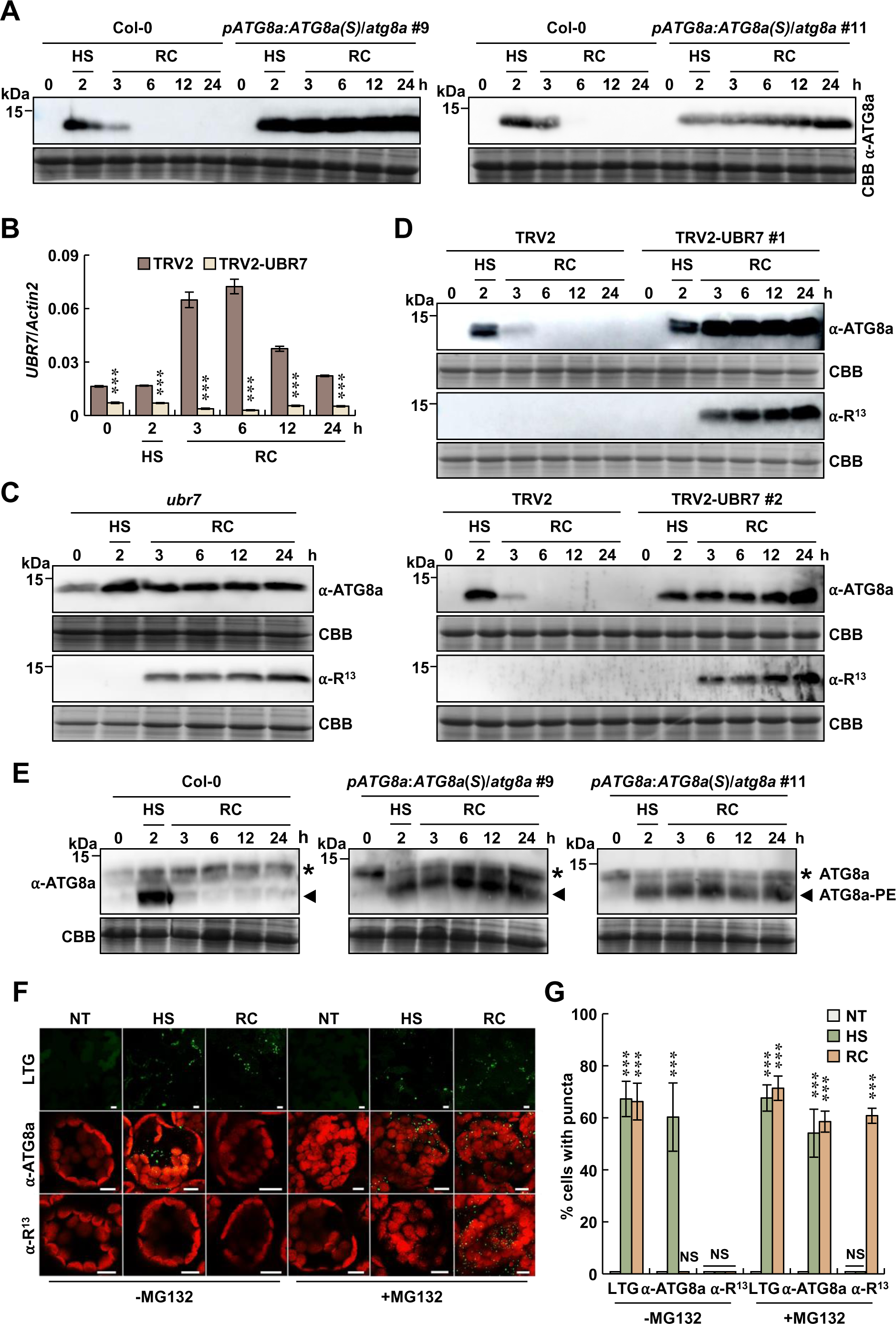
ATG8a degradation is UBR7-dependent during HS recovery. (A) Sustained ATG8(S) expression is tolerant to proteasomal degradation during HS recovery. ATG8a expression was monitored during HS and recovery in *pATG8:ATG8a*(*S*)/*atg8a* plants. (B) *UBR7* is highly induced during HS recovery but suppressed by VIGS. *UBR7* transcript levels were analyzed by RT-qPCR in TRV2 control and TRV2-UBR7 plants exposed to HS (42°C) for 2 h and recovery (23°C) for the indicated times. Data represent means ± SD (*n* = 4 biological replicates). Asterisks indicate significant differences between TRV2 and TRV2-UBR7 plants (*t* test; \*\*\**P* < 0.001). (C and D) ATG8a degradation is UBR7-dependent during HS recovery. ATG8a expression was monitored during HS and recovery in *ubr7* (C) and TRV2 and TRV2-UBR7 (D) plants. α-R^13^, anti-R^13^-ATG8a antibody. (E) ATG8a lipidation assay. Protein extracts were prepared from Col-0 and *pATG8:ATG8a*(*S*)/*atg8a* plants exposed to HS (42°C) for 2 h and recovery (23°C) for the indicated times. Proteins were separated by SDS-PAGE in the presence of 6 M urea. Asterisks and arrowheads indicate ATG8a and ATG8a-PE, respectively. (F) LTG staining and immunostaining with the anti-ATG8a and anti-R^13^-ATG8a antibodies for autophagic structures in Col-0 exposed to HS (42°C) for 2 h and recovery (23°C) for 6 h. For MG132 treatment, plants were sprayed with MG132 (10 μM) prior to exposure to HS and recovery conditions. Bars, 10 μm. (G) Quantification of cells with LTG-stained and ATG8a/R^13^-ATG8a-positive autophagic puncta in (F). Stained leaves were photographed and cells with fluorescent spots were counted per 250 μm^2^ area. Results represent means ± SD (*n* = 6). Asterisks indicate significant differences from the respective NT (*t* test; \*\*\**P* < 0.001). NS, not significant. ATG8a proteins were analyzed by immunoblotting with respective antibodies, and Coomassie brilliant blue (CBB) staining served as a loading control. NT, no treatment; HS, heat stress; RC, recovery.

Next, we assessed autophagy activation during HS and recovery. ATG8a expression in wild-type and *pATG8:ATG8a*(*S*)/*atg8a* plants correlated with the formation of active ATG8a, i.e., the ATG8a-PE conjugate (Figure 5E). Moreover, the formation of autophagosomes, a hallmark of autophagy activation, was monitored in wild-type plants by LysoTracker Green (LTG) staining (Kwon et al., 2013) and immunostaining with the anti-ATG8a and R^13^-ATG8a antibodies (Figures 5F and 5G). While LTG-stained punctate structures highly accumulated during both HS and recovery, ATG8a-positive autophagic structures were detected only under HS. MG132 treatment restored the accumulation of ATG8a- and R^13^-ATG8a-positive puncta during HS recovery, consistent with the expression patterns of ATG8a- and R^13^-ATG8a (Figure 4D). These results suggest that while ATG8a functions in HS-induced autophagy, other ATG8 isoforms may replace ATG8a to participate in autophagy during HS recovery. Thus, we analyzed the expression of other *ATG8s* during HS responses, especially whether their expression increases after HS release. Supporting our idea, *ATG8b*, *ATG8d*, and *ATG8g* were significantly induced during HS recovery (Figure S5A).

### ATG8a turnover is required for tolerance to recurring HS in *Arabidopsis*

To assess the role of ATG8a in HS responses, we additionally constructed transgenic lines of *atg8a* complemented with *ATG8a* genomic DNAs containing either the wild-type sequence (*gATG8a*/*atg8a*) or a G to A mutation at the 5’ splice site of the retained intron in ATG8a(L) (*gATG8a^PM^*/*atg8a*), which exclusively produces *ATG8a*(*L*) (Figure S5B). We then examined how these genomic DNA complementation lines express ATG8a during HS and recovery. As expected, *gATG8a*/*atg8a* lines exhibited the same ATG8a expression patterns as wild-type plants at both RNA and protein levels (Figures S5C and S5D). However, *gATG8a^PM^*/*atg8a* lines failed to express *ATG8a*(*S*) (Figure S5C) and barely produced ATG8a proteins under HS, but expressed degradable ATG8a(L) during HS recovery (Figure S5D). Under HS, the aberrantly generated *ATG8a(L)* transcripts, due to the failure of intron removal, appear to be destabilized in *gATG8a^PM^*/*atg8a* lines.

In nature, temperature fluctuates during the day and night, exposing plants to repeated daily HS and recovery, particularly during hot summer days. To examine ATG8a expression under such conditions, plants were grown at 23 °C under long-day conditions (16-h light/8-h dark cycle) with 12-h HS at 42°C in the middle of the day. Notably, the increase and decrease in ATG8a levels during HS and recovery were repeated daily (Figure 6A). Recurring ATG8a turnover was also evident under short-day conditions (8-h light/16-h dark cycle) with 8-h HS during the day (Figure S6A). ATG8a expression was not detectable during the day and night under normal conditions, indicating that daily changes in ATG8a abundance do not correlate with circadian rhythm.

**Figure 6.**
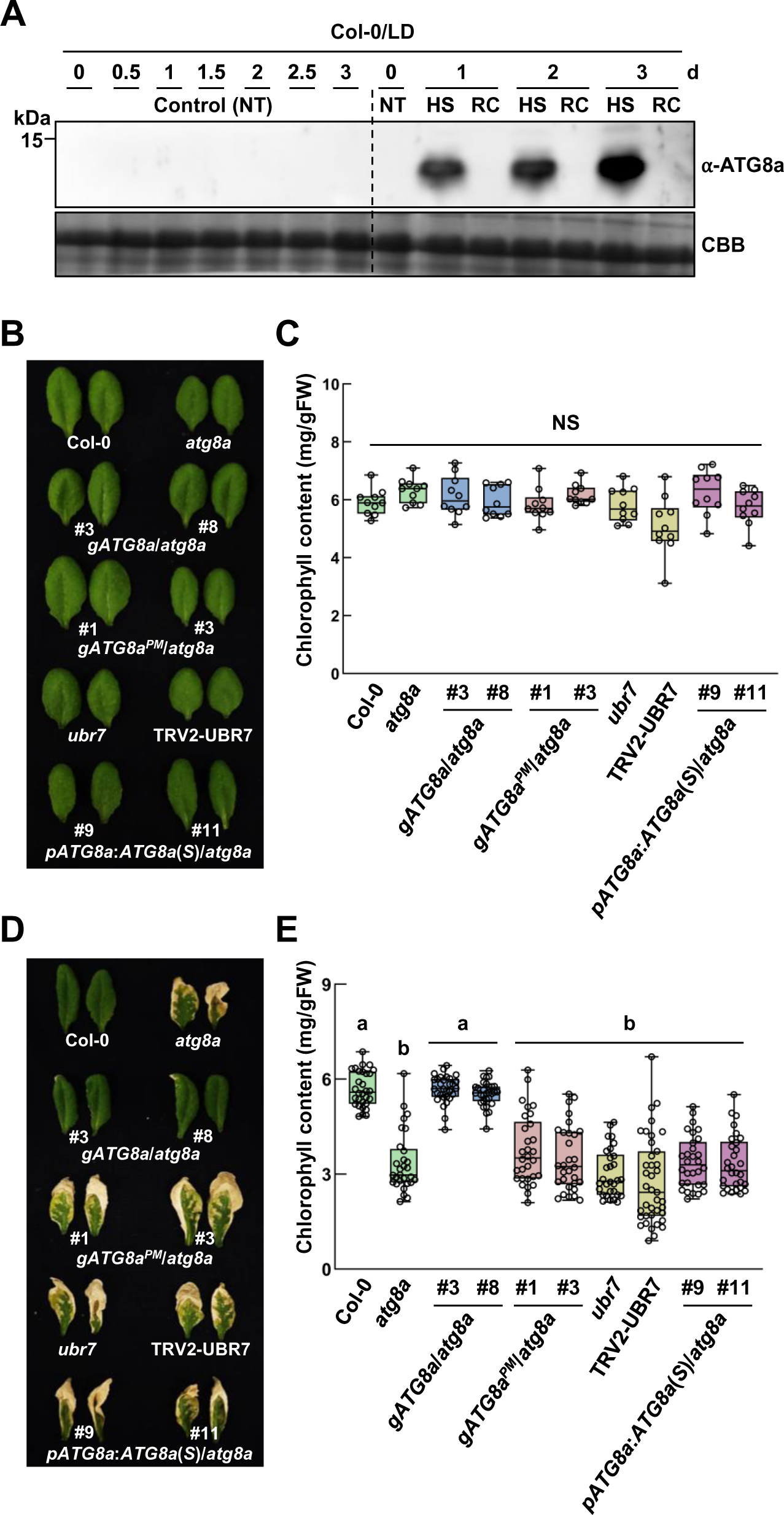
ATG8a turnover is important for plant thermotolerance. (A) Fluctuations in ATG8a abundance coincide with daily HS and recovery cycles. Col-0 plants grown in long days were exposed to 42°C (HS) for 12 h during the day and maintained at 23°C (recovery) for the remaining 12 h. Total proteins were extracted from Col-0 plants exposed to HS and recovery repeatedly for 3 days and analyzed by immunoblotting with the anti-ATG8a antibody. Coomassie brilliant blue (CBB) staining served as a loading control. LD, long day; NT, no treatment; HS, heat stress; RC, recovery. (B) Leaf phenotypes of Col-0, *atg8a*, *ubr7*, TRV2-UBR7, and complementation lines grown in long days. Plants were maintained at 23°C under long-day conditions during HS experiments. (C) Quantification of chlorophyll content in control plants in (B). Data represent means ± SD (*n* = 10 leaves). NS, not significant. (D) Thermotolerance phenotypes of Col-0, *atg8a*, *ubr7*, TRV2-UBR7, and complementation lines under recurring HS and recovery conditions. Plants grown in long days were exposed to 42°C (HS) for 12 h during the day and maintained at 23°C (recovery) for the remaining 12 h, which was repeated for 5 days. Plants were allowed to recover for an additional 3 days and observed for leaf phenotypes. (E) Quantification of chlorophyll content in plants exposed to HS and recovery in (D). Data represent means ± SD (*n* > 30 leaves). Different letters indicate significant differences (Tukey’s HSD test; *P* < 0.05).

Next, *atg8a*, *ubr7*, TRV2-UBR7, and all complementation lines were subjected to repeated daily HS and recovery for 5 days and evaluated for thermotolerance. Based on the ATG8a levels, they can be divided into three classes: (1) *atg8a* null mutant, (2) *gATG8a^PM^*/*atg8a* lines with defective ATG8a under HS (Figure S5D), and (3) *ubr7*, TRV2-UBR7, and *pATG8:ATG8a*(*S*)/*atg8a* lines with high ATG8a expression (Figures 5A, 5C, and 5D). Unlike wild-type and *gATG8a*/*atg8a* plants, which remained green, all other tested plants exhibited reduced thermotolerance under both long-day and short-day conditions; their leaves turned yellow with a significant decrease in chlorophyll content (Figures 6B-6E and S6B-S6E). These results suggest that daily ATG8a turnover is important for plant response to recurring HS and recovery, and that abnormally low or excessive ATG8a expression both impair plant thermotolerance.

### Proteomic analysis reveals ATG8a-dependent altered abundance of HS-responsive proteins during HS recovery

Previous studies report that autophagy is critical for degradation of thermo-sensitive proteins and HSPs during both the HS and recovery periods (Zhou et al., 2013; Sedaghatmehr et al., 2019, 2021; Thirumalaikumar et al., 2021). In this study, we showed that HS-upregulated ATG8a abundance decreases and other *ATG8s* such as *ATG8b*, *ATG8d*, and *ATG8g* are activated after release from HS (Figures 4C, 4D, and S5A). This implies that other ATG8 isoforms may replace ATG8a to induce autophagic protein degradation during HS recovery. The results of previous and our studies led us to hypothesize that autophagic cargo to be cleared may change at different stages of HS responses and be recognized by ATG8a and other ATG8s that are differentially expressed during the HS and recovery periods.

Based on our speculation that ATG8 isoforms differentially target autophagic cargo during distinct stages of HS responses, we aimed to identify proteins whose clearance is disturbed by sustained ATG8a expression during HS recovery. For this, we analyzed proteomes of wild-type and *pATG8:ATG8a*(*S*)/*atg8a* (#9) plants subjected to HS and recovery using label-free liquid chromatography-tandem mass spectrometry (LC-MS/MS) (Table S1). Significant changes in protein expression were defined according to the cutoff criteria (adjusted *P* < 0.05, log_2_ fold change (|log_2_FC)| ≥ 1). HS-responsive proteins were first identified by analyzing differentially expressed proteins (DEPs) between control (no treatment, NT) and HS treatment, revealing 149 (93 increased and 56 decreased) and 133 (85 increased and 48 decreased) DEPs (HS-DEPs) in wild-type and *pATG8:ATG8a*(*S*)/*atg8a* plants, respectively (Figures 7A and 7B; Table S2). All HS-DEPs in *pATG8:ATG8a*(*S*)/*atg8a* plants overlapped entirely with those in wild-type plants, meaning that the HS response of *pATG8:ATG8a*(*S*)/*atg8a* plants is essentially the same as that of wild-type plants. The upregulated HS-DEPs were then subjected to Gene Ontology (GO) enrichment analysis using GO Biological Process (BP) terms provided by the PANTHER database (http://pantherdb.org) (Table S3). The top significant GO terms indicated that the upregulated HS-DEPs in wild-type and *pATG8:ATG8a*(*S*)/*atg8a* plants are commonly involved in biological processes such as heat response and protein folding (Figure 7C). These results further suggest that ATG8a(S) acts as a major ATG8 in HS-activated autophagy.

**Figure 7.**
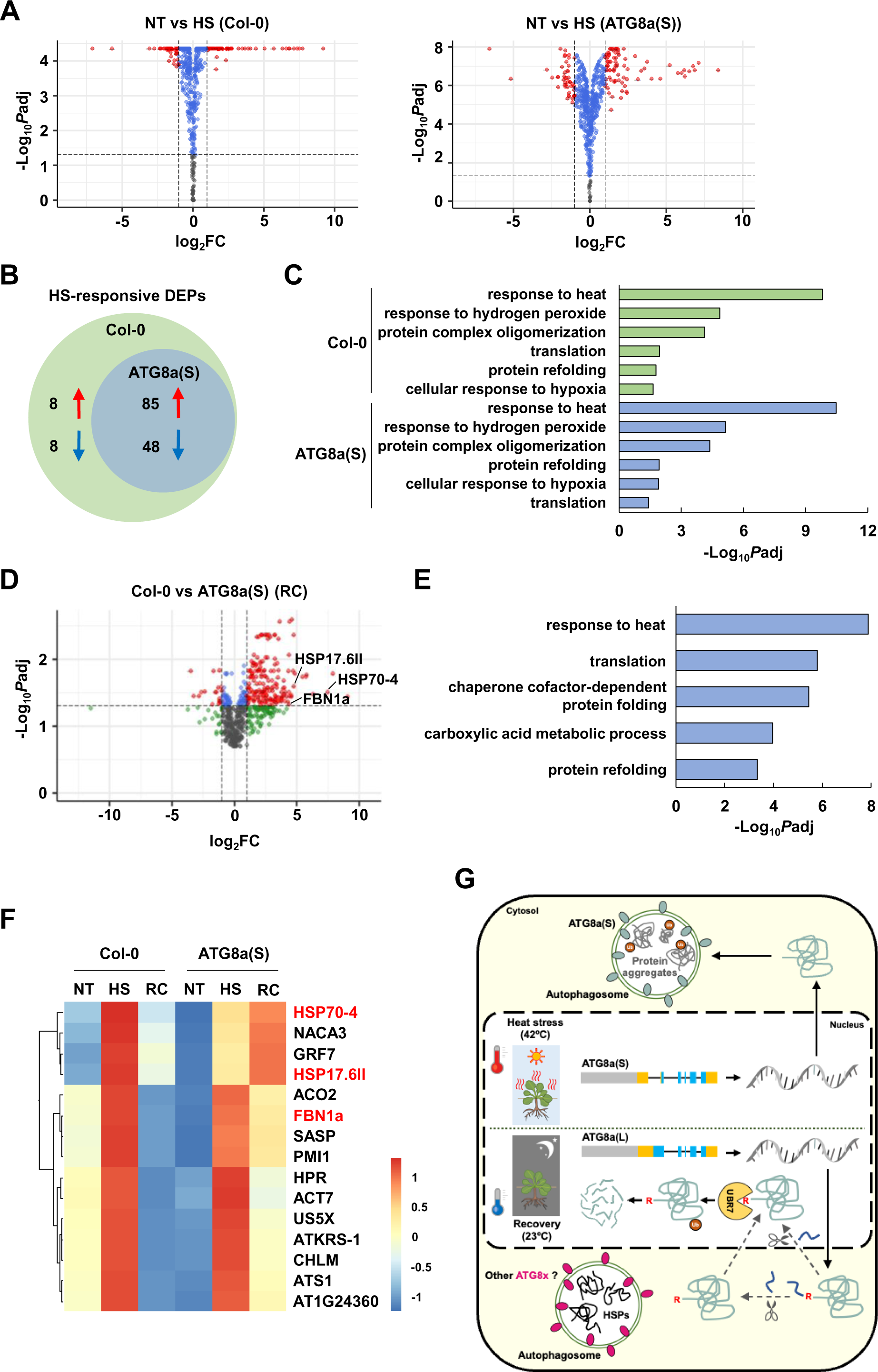
Proteomics reveals that ATG8a expression alters protein profiles during HS recovery. (A) Volcano plots of DEPs between no treatment (NT) and HS in Col-0 and *pATG8:ATG8a*(*S*)/*atg8a* plants. Cutoff values (*P*adj = 0.05 and |log_2_FC| = 1) are indicated by dashed lines. Red dots represent significantly upregulated and downregulated DEPs. Volcano plots were generated using the EnhancedVolcano package in R Studio. (B) Venn diagrams for HS-responsive DEPs in Col-0 and *pATG8:ATG8a*(*S*)/*atg8a* plants. (C) GO enrichment analysis of HS-upregulated DEPs in Col-0 and *pATG8:ATG8a*(*S*)/*atg8a* plants. The 6 most significantly (FDR < 0.05) enriched GO terms in the Biological Process are presented for HS-upregulated DEPs. (D) Volcano plots of DEPs between Col-0 and *pATG8:ATG8a*(*S*)/*atg8a* plants during HS recovery (RC). Cutoff values (*P*adj = 0.05 and |log_2_FC| = 1) are indicated by dashed lines. Red dots represent significantly upregulated and downregulated DEPs. Volcano plots were generated using the EnhancedVolcano package in R Studio. (E) GO enrichment analysis of DEPs between Col-0 and *pATG8:ATG8a*(*S*)/*atg8a* plants during RC. The 5 most significantly (FDR < 0.05) enriched GO terms in the Biological Process are presented. (F) Heatmap showing expression profiles of 15 selected DEPs between Col-0 and *pATG8:ATG8a*(*S*)/*atg8a* plants under NT, HS, and RC. Selected proteins were HS-upregulated DEPs that decreased in Col-0 but remained high in *pATG8:ATG8a*(*S*)/*atg8a* plants during RC. Protein information is provided in Table S6. Heatmap was generated using the pheatmap package in R Studio. (G) *ATG8a* pre-mRNAs are alternatively spliced to generate two splice variants *ATG8*(*S*) and *ATG8a*(*L*), which accumulate mainly during HS and recovery, respectively. ATG8a(L) is N-terminally processed to expose the Arg/N-degron, which is recognized by the N-recognin UBR7, leading to polyubiquitination and subsequent proteasomal degradation of ATG8a. Eventually, ATG8a and other ATG8 isoforms (ATG8x) may participate in selective autophagy, targeting different cargos, e.g., protein aggregates and HSPs, during HS and recovery phases, respectively. These fluctuations in ATG8a abundance coincide with daily thermocycles, enhancing thermotolerance in plants. FC, fold change; NT, no treatment; HS, heat stress; RC, recovery; ATG8a(S), *pATG8:ATG8a*(*S*)/*atg8a* plants.

Next, we analyzed proteins (RC-DEPs) that are differentially expressed between wild-type and *pATG8:ATG8a*(*S*)/*atg8a* plants during HS recovery (Table S4). Of the identified RC-DEPs, the majority (183 of 202, 91%) were up-regulated in *pATG8:ATG8a*(*S*)/*atg8a* plants, as displayed in the volcano plot (Figure 7D). The GO analysis of RC-DEPs revealed that 183 up- regulated RC-DEPs are enriched in BP terms of heat response and protein folding (Figure 7E; Table S5), similar to the upregulated HS-DEPs. A heatmap was generated to show the changes in protein abundance among NT, HS, and recovery conditions (Figure S7). The proteomic profiles revealed that 15 upregulated HS-DEPs are decreased in wild-type but remain upregulated in *pATG8:ATG8a*(*S*)/*atg8a* plants during HS recovery (Figure 7F; Table S6). Among them, two HSPs, HSP70-4 and HSP17.6II, and fibrillin FBN1a (Gámez-Arjona et al., 2014; Espinoza-Corral et al., 2021) were annotated to the GO term for response to heat. Together, these results support our hypothesis that sustained ATG8a expression may interfere with the clearance of HS-induced proteins, including HSPs, which would otherwise disappear in wild-type plants during HS recovery.

## DISCUSSION

In this study, we demonstrate that ATG8a stability is regulated by the Arg/N-degron pathway, resulting in the removal of ATG8a proteins. *ATG8a* pre-mRNAs are alternatively spliced to produce two splice variants *ATG8*(*S*) and *ATG8a*(*L*), with *ATG8a*(*L*) being an IR variant encoding N-degron-bearing ATG8a. *ATG8*(*S*) and *ATG8a*(*L*) exhibit differential expression patterns, coinciding with timely fluctuations in ATG8a abundance. We propose a molecular mechanism for plant adaptation to HS via N-degron-mediated proteolytic regulation ATG8a (Figure 7G).

Our results indicate that AS is crucial for dynamic changes in ATG8a abundance during HS responses (Figures 4A-4C). AS is a post-transcriptional regulatory process that enhances the functional diversity of proteins and plays an import role in plant responses to abiotic stress (Chaudhary et al., 2019). Among AS events, IR is the most frequent one in plants, while exon skipping is prevalent in mammals (Reddy et al., 2013). Many studies have addressed the role of AS in plant HS responses through the generation of active splice isoforms of heat-responsive regulators, including heat shock transcription factors (HSFs) (Laloum et al., 2018; Ling et al., 2021). Our finding makes an important contribution to these studies, demonstrating that AS provides a means of fine-tuning gene expression in response to HS. We also found that AS generates different 5’ UTRs in *ATG8*(*S*) and *ATG8a*(*L*) variants, with a specific sequence (designated as L) in *ATG8a*(*L*) 5’ UTR enhancing translational efficiency. This is a plausible mechanism for the increased production of ATG8a(L) proteins during the post-HS period (Figures 4E-4H). Further investigations are required to identify *cis*-RNA elements and *trans*-factors that promote translation of *ATG8a*(*L*) transcripts.

UBR7 was previously identified as a histone H2B monoUb ligase, which suppresses breast tumors (Adhikary et al., 2019). In addition, TurboID-based proximity labeling identified UBR7 as a regulator of the tobacco N immune receptor (Zhang et al., 2019). Our results provide the first line of evidence that UBR7 acts as an N-recognin and, like other UBRs, recognizes ATG8a(L) with an N-terminal Arg through the UBR box domain. Notably, while ATG8(L) proteins localize in both the cytosol and nucleus, UBR7 is expressed and interacts with ATG8a(L) in the nucleus (Figures 3D, S2G, and S3), suggesting that N-degron-mediated ATG8a degradation occurs in the nucleus. Considering that ATG8s are synthesized as cytosolic factors and are involved in autophagosome biogenesis, spatial separation of ATG8a activation and degradation into the cytosol and nucleus may be necessary to avoid the collision of two contradictory events. We also show that N-terminal cleavage of ATG8a(L) is a prerequisite for UBR7-mediated polyubiquitination and degradation (Figure 2). A number of N-degron substrates are created through decapping (Gibbs et al., 2011; Licausi et al., 2011; Gibbs et al., 2014a; Vicente et al., 2017; Gibbs et al., 2018; Dissmeyer, 2019) or endoproteolytic cleavage (Piatkov et al., 2012; Xu et al., 2012; Brower et al., 2013; Dong et al., 2017; Chui et al., 2019; Dissmeyer, 2019; Sandstrom et al., 2019; Xu et al., 2019; Chen et al., 2021). Given that the *Arabidopsis* genome contains over 700 genes encoding putative proteases (Garcia-Lorenzo et al., 2006), the specific protease(s) for endoproteolytic processing of ATG8a(L) remains to be determined.

Autophagy provides the most effective way to remove protein aggregates and damaged organelles and plays a homeostatic role in plants exposed to abiotic stress, including HS (Zhou et al., 2013). We propose that ATG8a degradation may be necessary to replace ATG8a with other ATG8s (e.g., ATG8b, ATG8d, and ATG8g) (Figure S5A), enabling distinct ATG8s and their interacting cargo receptors to selectively target different cargo in the process of selective autophagy during HS and recovery (Figure 7G). Previous studies have shown that autophagy receptors determine the specificity of cargo, and their interactions with ATG8 isoforms vary depending on cargo and biological processes involved (Luo et al., 2021). The proposed mechanism is supported by our proteomics results, which show that HS-upregulated proteins decrease in wild-type plants but remain high in *pATG8:ATG8a*(*S*)/*atg8a* plants during the HS recovery phase (Figures 7D and 7F). We speculate that sustained ATG8a expression interferes with the actions of other ATG8(s) in *pATG8:ATG8a*(*S*)/*atg8a* plants, thereby preventing degradation of heat-responsive proteins, including HSPs, which would otherwise decrease in abundance in wild-type plants released from HS. Our findings are consistent with previous studies, in which autophagy is directed at distinct targets during HS and recovery, mediating selective degradation of HSPs during the recovery phase (Sedaghatmehr et al., 2019; Thirumalaikumar et al., 2021). The removal of heat-responsive proteins during HS recovery may be required to reset stress conditions and enhance resistance to upcoming HS. A recent study reveals a non-canonical role of ATG8s in Golgi reassembly after HS (Zhou et al., 2023). Whether proteolytic regulation of ATG8a is related to organellar recovery in plant response to HS remains to be explored.

In nature, plants are exposed to day and night temperature fluctuations, subjecting them to daily HS and recovery conditions. The increase and decrease in ATG8a abundance coincided with HS and recovery cycles, and abnormally low (*atg8a* and *gATG8a^PM^*/*atg8a* lines) and high (*ubr7*, TRV2-UBR7, and *pATG8:ATG8a*(*S*)/*atg8a* lines) levels of ATG8 expression both impaired plant thermotolerance (Figures 6 and S6), implying that ATG8a fluctuations are critical for plant thermotolerance. Intriguingly, early studies on crops have reported that plants undergo a diurnal cycle of heat resistance, which peaks during the day and decreases at night (Laude, 1939). This is consistent with findings that a diurnal pattern of thermotolerance correlates with diurnal expression of *HSP* genes (Dickinson et al., 2018). Together, our findings demonstrate that the N-degron pathway is a critical mechanism for timely and quantitative regulation of ATG8a and, ultimately, heat-responsive proteins over daily thermocycles. This dynamic regulation may have evolved as a survival strategy for plants to cope with environmental thermal changes.

## METHODS

### Plant materials and growth conditions

*Arabidopsis thaliana* (Ecotype Columbia, Col-0) plants were grown at 23°C under long-day conditions in a 16-h light/8-h dark cycle and short-day conditions in an 8-h light/16-h dark cycle. The mutant lines used in this study are *prt6-1* (SALK_004079), *big-3* (SALK_107817) (Bruggeman et al., 2020), *ubr7* (SALK_034619), and *atg8a* (SALK_045344) from the Arabidopsis Biological Resource Center (ABRC). T-DNA insertion sites were verified by sequence analysis using gene-specific primers (Table S7). To generate *pATG8:ATG8a*(*S*)/*atg8a* plants, the *ATG8a*(*S*) coding region was cloned into the pCAMBIA3300 binary vector under the control of the native p*ATG8a* promoter (-1 to -1150 bp relative to the transcription start site) amplified from *Arabidopsis* genomic DNA by PCR. To generate *gATG8a*/*atg8a* and *gATG8a^PM^*/*atg8a* plants, the 2572-bp genomic DNA region containing *ATG8a* was amplified by PCR, and a G to A mutation was made by site-directed mutagenesis using primers in Table S7. The constructs were transformed into *atg8a* plants via *Agrobacterium tumefaciens* GV3101 using the floral dip method (Clough and Bent, 1998).

### Plant treatments

For chemical treatments, *Arabidopsis* protoplasts were treated with MG132 (10 µM with 0.1% DMSO) and cycloheximide (100 µM with 0.1% DMSO) for 3h before sampling. Four-week-old plants were sprayed with MG132 (10 µM with 0.1% DMSO) or syringe-infiltrated with DTT (8 mM in H_2_O) and TM (250 ng/ml with 0.1% DMSO) and incubated for the indicated times. HS experiments were performed as previously described (Silva-Correia et al., 2014). For expression analysis, 4-week-old plants were exposed to 42°C (HS phase) and then incubated at 23°C (recovery phase) for the indicated times in an incubator. For phenotype and chlorophyll content analysis, HS treatment at 42°C was performed for 8 or 12 h during the day on 4-week-old plants grown under short-day or long-day conditions, respectively, which was repeated for 5 days. Plants were allowed to recover for additional 3 days before further analysis.

### Gene expression analysis

Total RNAs were extracted from *Arabidopsis* leaves using TRIzol reagent (Meridian Bioscience), treated with DNase I (New England Biolabs), and reverse-transcribed into cDNAs using PrimeScript RT reagent kit (TaKaRa). Quantitative real-time PCR (RT-qPCR) was performed using KAPA SYBER FAST qPCR master mix (Kapa Biosystems) with gene-specific primers (Table S7) on a LightCycler 480 system (Roche) according to the manufacturer’s protocol. *Actin2* was used as the reference gene for normalization. Data were analyzed using LC480 Conversion and LinRegPCR software (Heart Failure Research Center).

### Transient expression assay

Various expression constructs were generated for transient expression assays in *Arabidopsis* protoplasts. For protein expression constructs, coding regions of *ATG8a*, *ATG8e*, *UBR7*, and *Ub* were amplified from the *Arabidopsis* cDNA library by PCR and cloned into pUC19 vectors containing CaMV 35S promoter, *RBCS* terminator, and the indicated tag sequences. Deletion and substitution mutations were introduced into *ATG8a* by site-directed mutagenesis using primers in Table S7. For 5’ UTR-driven reporter (translational efficiency) constructs, the *GFP* sequence was fused with the indicated 5’ UTR sequences of *ATG8a*(*S*) and *ATG8a*(*L*) and the 3’ UTR sequence of *ATG8a,* and cloned into the pUC19 vector containing p*ATG8a* promoter and *RBCS* terminator sequences. Ub fusion constructs, Ub-R^13^-ATG8a, Ub-A^13^-ATG8a, Ub-R-GUS-FLAG, and Ub-M-GUS-FLAG, were generated through a DNA synthesis service at Bionics and cloned into pUC19 vectors containing CaMV 35S promoter and *RBCS* terminator.

*Arabidopsis* mesophyll protoplasts were isolated and transfected as previously described (Yoo et al., 2007). Leaves were sliced and incubated for 3 h in an enzyme solution (20 mM MES-KOH, pH 5.7, 1% cellulase R-10, 0.4% macerozyme R-10, 0.4 M D-mannitol, 20 mM KCl, 10 mM CaCl_2_, and 0.1% BSA). Protoplasts were collected by filtering through a 70-μm cell strainer, centrifuged, and resuspended in W5 solution (1.5 mM MES-KOH, pH 5.6, 154 mM NaCl, 125 mM CaCl_2_, and 5 mM KCl). Isolated protoplasts (2 x 10^4^) were transfected with 15 μg of *ATG8a* and *ATG8e*, 20 μg of *UBR7*, 10 μg of *Ub*, 15 μg of 5’ UTR reporters, 20 μg of BiFC constructs, and 10 μg of Ub fusion constructs. After transfection, protoplasts were incubated at room temperature overnight to allow expression.

### Antibody preparation

A rabbit polyclonal antibody specific for R^13^-ATG8a was generated using the peptide RIAMAKSSFKI corresponding to the N-terminal sequence (positions 13-23) of ATG8a through a custom service at AbFrontier as previously described (Cha-Molstad et al., 2015). Briefly, antisera from rabbits immunized with the peptide were subjected to affinity purification using the peptide RIAMAKSSFKI. The specificity of the purified antibody was validated by immunoblot analysis using R^13^-ATG8a and A^13^-ATG8a and the peptide competition assay.

### Immunoblotting and co-immunoprecipitation

For immunoblotting, total proteins were extracted by boiling protoplasts and ground plant leaves with liquid nitrogen in 6x sample buffer (100 mM Tris-HCl, pH 6.8, 12% SDS, 20% glycerol, 6% β-mercaptoethanol, and 0.2% bromophenol blue) for 10 min. For co-immunoprecipitation, protoplasts were lysed in lysis buffer (50 mM Tris-HCl, pH 7.4, 150 mM NaCl, 1% NP-40, and 1x protease inhibitor cocktail) for 30 min on ice. Lysates were centrifuged at 13,000 rpm for 10 min at 4°C, and the supernatant was incubated with the anti-HA antibody (Thermo Fisher Scientific) for 1 h at 4°C. After an additional overnight incubation with Protein A agarose beads (Pierce), beads were washed with lysis buffer, and bound proteins were eluted by boiling in 6x sample buffer. Proteins were separated by SDS-PAGE and transferred to polyvinylidene fluoride membranes. Membranes were incubated with anti-HA (Thermo Fisher Scientific, 71-5500), anti-MYC (Abcam, ab32), anti-ATG8a (Abcam, ab77003), anti-Ub (Santa Cruz, sc-8017), anti-FLAG (Abcam, ab205606), anti-mCherry (Abcam, ab183628), and anti-GFP (Santa Cruz, sc-9996) antibodies. Antibody-bound proteins were detected by incubation with secondary antibodies conjugated to horseradish peroxidase using an enhanced chemiluminescence system (Amersham Biosciences).

### Protein expression and purification

The coding regions of *ATG8a-i* were cloned into the modified pMAL2 vector containing the 6xHis and *FLAG* sequences to generate ATG8 fused to the N-terminal His and FLAG tags. The coding region of *UBR7* was cloned into the modified pMAL2 vector containing the *MYC* sequence to generate UBR7 fused to the N-terminal MYC tag. Substitution mutations for UBR7^CM^ were introduced into *UBR7* by site-directed mutagenesis using primers in Table S7. *UBC8* encoding the *Arabidopsis* E2 enzyme was cloned into the pMAL2 vector to generate UBC8 fused to the N-terminal maltose-binding protein (MBP) tag. *Escherichia coli* BL21(DE3) pLysS cells were transformed with the constructs and cultured at 37°C. Protein expression was induced by the addition of 0.3 mM isopropyl-β-D-thiogalactoside (IPTG) for 3 h at 28°C. The cells were harvested and lysed by sonication in lysis buffer (50 mM Tris-HCl, pH 7.4, 400 mM NaCl, 5% glycerol, 1 mM MgCl_2_, 5 mM Imidazole, 2 mM DTT, 0.5 mM PMSF, 1 mM NaF, 1 mM Na_3_VO_4_, and 1x protease inhibitor cocktail). His-FLAG-ATG8 proteins were purified using Ni-NTA agarose (QIAGEN), and MYC-UBR7 and MBP-UBC8 proteins were purified using anti-MYC (Pierce) and anti-MBP (New England Biolabs) magnetic beads, respectively, according to the manufacturer’s instructions.

R^13^-ATG8a and A^13^-ATG8a proteins were prepared as previously described (Kim et al., 2020). ATG8a was cloned into the modified pET vector to generate GST-LC3B-R^13^/A^13^-ATG8a fusion proteins. *E. coli* BL21(DE3) cells were transformed with the constructs and cultured at 37°C. Protein expression was induced by the addition of 0.5 mM IPTG for 20 h at 18°C. The cells were harvested and lysed by sonication in PBS buffer (137 mM NaCl, 2.7 mM KCl, 10 mM Na_2_HPO_4_, and 1.8 mM KH_2_PO_4_). The expressed proteins were purified using a Glutathione Sepharose HP column (Cytiva). The eluted proteins were loaded onto a HiTrap Q HP column (Cytiva) and eluted with a NaCl gradient up to 1 M. The GST-LC3B tag was cleaved with human ATB4B protease (Kwon et al., 2017) overnight at 20℃ and eliminated using a Glutathione Sepharose HP column (Cytiva). R^13^/A^13^-ATG8a proteins were further purified by size-exclusion chromatography using a HiLoad 16/600 Superdex 75 pg column (Cytiva).

### *In vitro* pull-down assay

MYC-UBR7 (1 μg) was incubated with R^13^/A^13^-ATG8a (200 ng) in binding buffer (50 mM Tris-HCl, pH 7.4, 5 mM MgCl_2_, and 30% glycerol) for 2 h at 4°C. Anti-MYC antibody (Abcam, ab32) and protein G agarose beads (Pierce) were added and incubated for 2 h at 4℃. Bound proteins were eluted by boiling in 6x sample buffer, separated by SDS-PAGE, and visualized by immunoblotting with anti-ATG8a (Abcam, ab77003) and anti-MYC (Abcam, ab32) antibodies.

### *In vitro* ubiquitination assay

*In vitro* ubiquitination assay was performed as previously described (Wang et al., 2019). Recombinant MYC-UBR or MYC-UBR7^CM^ (500 ng) was incubated with either R^13^-ATG8a or A^13^-ATG8a (1 µg) in the presence of Ub (15 µg), human E1 UBE1 (50 ng; Merck), *Arabidopsis* E2 MBP-UBC8 (200 ng) in ubiquitination buffer (50 mM Tris-HCl, pH 7.4, 5 mM MgCl_2_, 2 mM DTT, 4 mM ATP, and 1x protease inhibitor cocktail) for 3 h at 30°C. Reactions were stopped by boiling in 6x sample buffer, and proteins were separated on a gradient gel (Invitrogen) by SDS-PAGE and detected by immunoblotting with anti-ATG8a (Abcam, ab77003), anti-MYC (Abcam, ab32), and anti-Ub (Santa Cruz, sc-8017) antibodies.

### ATG8a lipidation assay

ATG8a lipidation assay was performed as previously described (Jeon et al., 2023). Total proteins were extracted from 4-week-old plants exposed to HS and recovery conditions. Proteins were separated on 15% gels containing 6 M urea by SDS-PAGE, transferred to polyvinylidene fluoride membranes (Amersham Biosciences), and detected by immunoblotting with the anti-ATG8a antibody (Abcam, ab77003).

### BiFC and subcellular localization

For BiFC, the full-length coding regions of *ATG8a*, *ATG8a* with the indicated mutations (*ATG8a^R13A^* and *ATG8a^3A^*) and *UBR7*, and the UBR domain region of *UBR7* were PCR-amplified using primers in Table S7. The PCR products were cloned into pUC-SPYNE and pUC-SPYCE vectors (Walter et al., 2004) for the expression of fusion proteins UBR7-GFP^N^, UBR-GFP^N^, ATG8a-GFP^C^, ATG8a^R13A^-GFP^C^, and ATG8a^3A^-GFP^C^. For subcellular localization, the PCR products of *UBR7* and *ATG8a* were cloned into pUC19 vectors containing the *GFP* and *mCherry* sequences for the expression of fusion proteins UBR7-GFP and ATG8a-mCherry. *Arabidopsis* protoplasts were transfected with these constructs and incubated for 20-24 h for protein expression. To visualize nuclei, protoplasts were treated with 3 µM DAPI (Merck) overnight. Fluorescent signals were observed using a confocal microscope (Zeiss LSM 700). Excitation and emission were set at 488/543 nm for GFP, 555/630 nm for mCherry, 639/710 nm for chlorophyll, and 350/465 nm for DAPI.

### LTG staining

LTG staining was performed as previously described (Kwon et al., 2013). Leaf discs were vacuum-infiltrated with 1 µM LTG DND-26 (Invitrogen) and incubated in the dark for 1 h. Images were acquired using a confocal microscope (Zeiss LSM 700) at 488/543 nm excitation and emission.

### Immunostaining

Immunostaining was performed as described previously (Kwon et al., 2013). Leaf discs were fixed in 4% paraformaldehyde and 10% dimethyl sulfoxide dissolved in PMEG buffer (50 mM PIPES, pH 6.9, 5 mM EGTA, and 5 mM MgSO_4_) for 3 h at room temperature. Samples were washed three times with PMEG buffer and incubated in PMEG buffer containing 0.5% pectolyase (Sigma), 0.5% Triton X-100, and 1% bovine serum albumin (BSA) for 30 min at 37°C. Samples were then washed six times with PMEG buffer and incubated with anti-ATG8a (Abcam, ab77003) and R^13^-ATG8a antibodies in 1x phosphate-buffered saline (PBS) containing 3% BSA overnight at 4°C. After washing with PBS, samples were incubated with fluorescein isothiocyanate-conjugated goat anti-rabbit IgG (AlexaFlour 488, Invitrogen) for 3 h at room temperature. Images were acquired using a confocal microscope (Zeiss LSM 700) at 488/543 nm excitation and emission.

### Virus-induced gene silencing

VIGS was performed as previously described (Yang et al., 2021). The *UBR7* sequence was amplified and cloned into the pTRV2 vector. The constructs were transformed into *Agrobacterium tumefaciens* GV3101. *A. tumefaciens* cells were cultured in LB media containing 10 mM MES-KOH (pH 5.7), 200 µM acetosyringone, 50 mg/l gentamycin, and 50 mg/l kanamycin overnight at 28°C, adjusted to an OD_600_ of 1.5, and infiltrated into the first two true leaves of 2-week-old plants. After 24-26 days, VIGS plants were subjected to HS and recovery tests and gene silencing was validated using gene-specific primers (Table S7).

### Chlorophyll quantification

To determine chlorophyll content, plant leaves were ground in liquid nitrogen and immersed in 80% (v/v) acetone to extract chlorophylls. After centrifugation at 3,000 rpm for 15 min, the supernatant was measured at 663 and 645 nm, and chlorophyll content was calculated using the formula as previously described (Lichtenthaler, 1987).

### Sample preparation for LC-MS/MS analysis

Protein samples were prepared for proteomic analysis as previously described (Wiśniewski et al., 2009). Total proteins were extracted from plant leaves using extraction buffer (100 mM Tris-HCl, pH 7.4, 12% SDS, 20% glycerol, and 1x protease inhibitor cocktail). Proteins were incubated with 5 mM tris(2-carboxyethyl)phosphine (TCEP) (Thermo Scientific) in 50 mM ammonium bicarbonate for 30 min at 37 °C. Proteins were then transferred to YM-10 Microcon filtration devices (Millipore) and centrifuged at 14,000 *g* for 15 min, followed by two successive washes with UA solution (8 M urea, 100 mM Tris–HCl, pH 8.5). 50 mM iodoacetamide in UA solution was added to the filter unit and incubated in the dark at room temperature for 45 min. After centrifugation at 14,000 *g* for 15 min, proteins were washed one time with UA solution and again three times with 50 mM ammonium bicarbonate. For tryptic digestion, proteins were incubated with Trypsin Gold (Promega) using a 1:50 ratio (w/w) of trypsin to proteins overnight at 37°C. Peptides were eluted by centrifugation at 14,000 *g* for 15 min, followed by two centrifugations with 50 mM ammonium bicarbonate and 0.1% (v/v) formic acid. Peptides were desalted using C18 microspin columns (Harvard Apparatus) and eluted with 0.1% (v/v) formic acid in 80% acetonitrile. Peptide samples were dried under vacuum and resuspended in a solution containing 2% (v/v) acetonitrile and 0.1% (v/v) formic acid. Peptide concentrations were measured using NanoDrop (Thermo Scientific).

### LC-MS/MS analysis

LC-MS/MS analysis was performed using a nanoElute LC system (Bruker Daltonics) coupled to the timsTOF Pro (Bruker Daltonics), using a CaptiveSpray nanoelectrospray ion source (Bruker Daltonics). Peptide digest (150 ng) was injected into a capillary C18 column (25 cm length, 75 µm inner diameter, 1.5 µm particle size; Bruker Daltonics). The mobile phases were 0.1% (v/v) formic acid in water (A) and 0.1% (v/v) formic acid in acetonitrile (B). Gradient elution with solvent A and B was carried out using the following settings: 120 min gradient of 2-17% B for 60 min, 17-25% B for 30 min, 25-37% B for 10 min, 37-80% B for 10 min, and 80% B for 10 min at a flow rate of 0.4 µl/min. Mass spectral data in the range of m/z 100-1700 were collected in PASEF mode (Meier et al., 2015). Ion mobility resolution was set to 0.60-1.60 V s/cm over 100 ms ramp time, which ramped the collisional energy stepwise. Ten PASEF MS/MS scans per cycle were used for data-dependent acquisition.

### Data analysis

After data acquisition, data files were processed using the Peaks Studio 10.5 software (Bioinformatics Solution) and matched to tryptic peptide fragments from the *A. thaliana* protein sequence database (UniprotKB *A. thaliana*, downloaded January 2023; 136447 entries) with up to three missed cleavages allowed (Zhang et al., 2012). The mass tolerance for precursor and fragment ions was set to 15 ppm and 0.05 Da, respectively. Variable modifications included methionine oxidation and N-terminal acetylation, and fixed modifications included carbamidomethylation of Cys residues. Significant peptide identifications were performed with false discovery rate (FDR) < 0.01.

Raw data were subjected to data preprocessing using R software. The DESeq2 package (Love et al., 2014) was used to normalize and transform the data. Specifically, the rlog transformation was applied to normalize the data and reduce the influence of unwanted technical variation. Additionally, batch effect correction was performed using the SVA package (Leek et al., 2012) to account for any systematic differences introduced by different experimental batches. The Mfuzz package (Kumar et al., 2007) was employed to cluster proteins based on their expression patterns over time. Results were visualized in R using EnhanchedVolcano (RRID:SCR 018931) and pheatmap (RRID:SCR 016418).

### Statistical analysis

Statistical analyses were performed using SPSS (Statistical Package for the Social Sciences). Significant differences between experimental groups were analyzed by one-way ANOVA with Tukey’s HSD test or unpaired Student’s *t* test for multiple or single comparisons, respectively. Detailed information about statistical analysis is provided in the figure legends. Statistical significance was set at *P* < 0.05. All experiments were repeated three to five times with similar results. For proteomics data, to compare the expression levels between groups, a moderated *t* test was performed using the limma package (Ritchie et al., 2015). This test calculates log fold changes and associated *p*-values for each protein, considering the variability within and between groups. To account for multiple testing, *p*-values were adjusted using appropriate methods, such as the Benjamini-Hochberg procedure. Proteins were considered differentially expressed significantly based on the following criteria: |log_2_FC| ≥ 1 and *P*adj < 0.05.

## Author contributions

O.K.P. conceived and directed the project. S.H.K. and O.K.P. designed the research. S.H.K. performed most of the experiments. J.S.P., M.H.L., and J.P. participated in immunoblotting and RT-qPCR analysis. J.S., J.K., K.Y., J.C., and J.B.S. conducted proteomics analysis. W.S.Y. and H.K.S. performed protein purification. S.H.K. and O.K.P. analyzed the data and wrote the manuscript. All authors contributed to the review and editing of the manuscript.

## Acknowledgements

This work was supported by a Korea University grant and National Research Foundation of Korea (NRF) grants (2018R1A5A1023599, RS-2024-00339452) from the Korean government (MSIP).

## Declaration of interests

The authors declare no competing interests.

## Supplemental figure legends

**Figure S1.**
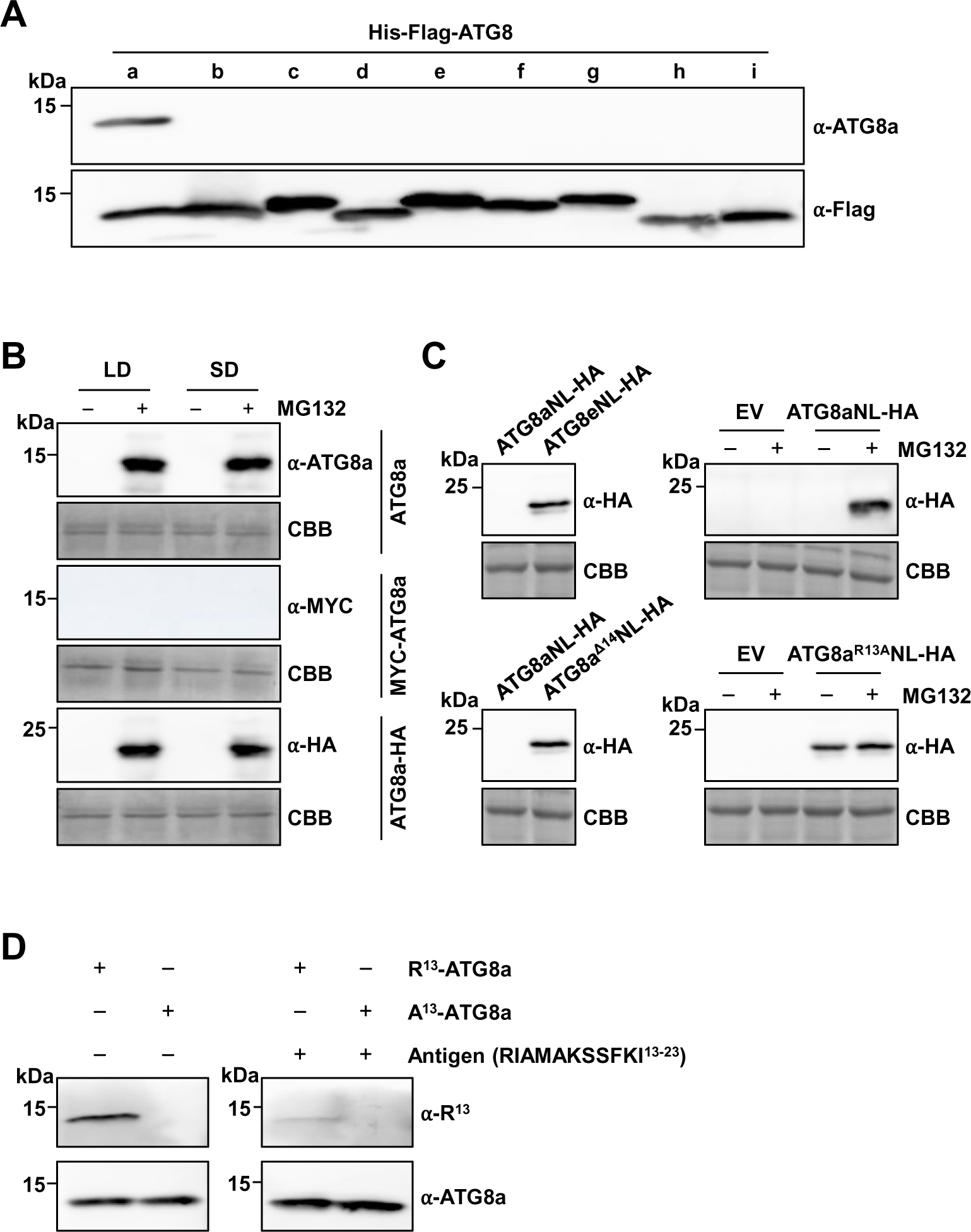
ATG8a undergoes 26S proteasome-dependent degradation independent of photoperiods and C-terminal cleavage, related to Figures 1 and 2. (A) Specificity test of the anti-ATG8a antibody. Recombinant proteins of His-Flag-ATG8 isoforms (ATG8a-i) were immunoblotted with the anti-ATG8a and anti-Flag antibodies. (B) ATG8a expression in different photoperiods. Untagged ATG8a, MYC-ATG8a, and ATG8a-HA were expressed in protoplasts prepared from Col-0 plants grown under long-day or short-day conditions, followed by treatments with cycloheximide (100 μM) and MG132 (10 μM) for 3 h. (C) Expression of non-lipidated (NL) ATG8a. ATG8aNL-HA, ATG8eNL-HA, ATG8a^Δ14^NL-HA, and ATG8a^R13A^NL-HA were expressed in Col-0 protoplasts, followed by treatments with cycloheximide (100 μM) and MG132 (10 μM) for 3 h. (D) Specificity test of the anti-R^13^-ATG8a antibody. Recombinant proteins of R^13^-ATG8a and A^13^-ATG8a prepared using the LC3B-fusion technique were immunoblotted with the anti-R^13^- ATG8a and anti-ATG8a antibodies in the presence or absence of the antigen peptide RIAMAKSSFKI at a 1:12 molar ratio of ATG8a to peptide. α-R^13^, anti-R^13^-ATG8a antibody. ATG8a proteins were analyzed by immunoblotting with respective antibodies, and Coomassie brilliant blue (CBB) staining served as a loading control.

**Figure S2.**
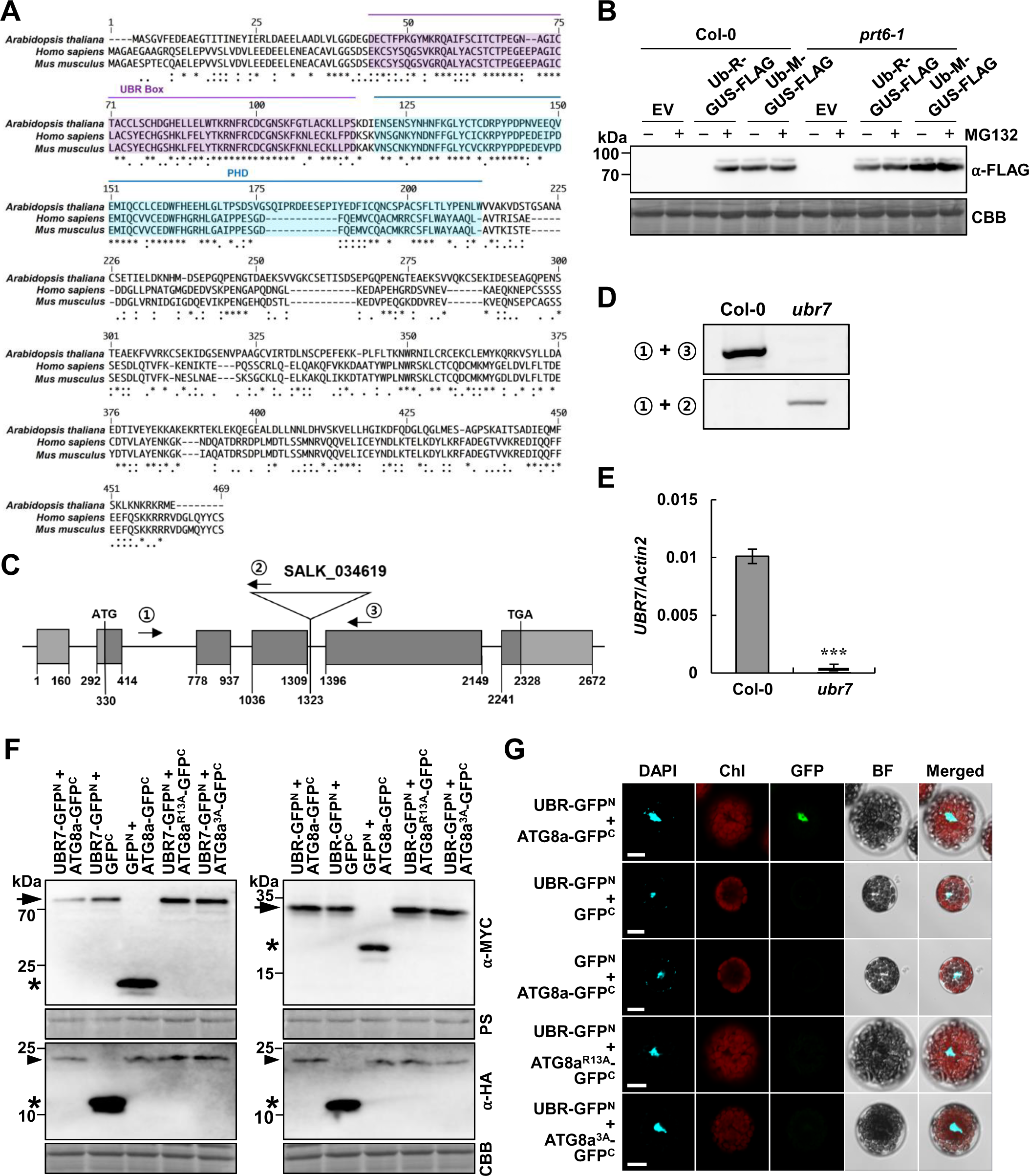
UBR7 interacts with ATG8a through the UBR box domain *in vivo*, related to Figure 3. (A) Multiple sequence alignment of UBR7 homologs in *Arabidopsis*, human, and mouse. Sequences were aligned using Clustal Omega online (https://www.ebi.ac.uk/Tools/msa/clustalo/). Conserved UBR and PHD domains are shaded in purple and blue, respectively. Consensus symbols are as follows: asterisks indicate identical residues; colons and periods indicate conserved and semiconserved substitutions, respectively. (B) R-GUS-FLAG generated through the Ub fusion technique is degraded in wild-type but stabilized in *prt6-1* mutant. Ub-R/M-GUS-FLAG were expressed in Col-0 and *prt6-1* protoplasts, followed by treatments with cycloheximide (100 μM) and MG132 (10 μM) for 3 h. GUS proteins were analyzed by immuniblotting with the anti-FLAG antibody, and Coomassie brilliant blue (CBB) staining served as a loading control. EV, empty vector. (C) Genomic structure of *UBR7* gene. Triangle and arrows indicate the positions of the T-DNA insertion and primers used for PCR, respectively. Genomic DNA sequences are represented by exons (dark gray boxes), introns (lines), and UTRs (light gray boxes). Numbers refer to nucleotides of *UBR7* gene. (D) Genotyping of *ubr7* mutant. PCR using primers indicated in (C) verified T-DNA insertion. (E) RT-qPCR analysis of *UBR7* expression in Col-0 and *ubr7* plants. *Actin2* was used as a control. Data represent means ± SD (*n* = 4 biological replicates). Asterisks indicate significant differences between Col-0 and *ubr7* plants (*t* test; \*\*\**P* < 0.001). (F) Validation of expression of GFP^N^-fused UBR7 (left) and UBR box (right) and GFP^C^-fused ATG8a, ATG8a^R13A^, and ATG8a^3A^ in transfected protoplasts. Protein lysates were prepared from Col-0 protoplasts transfected with the indicated constructs, followed by treatment with MG132 (10 μM) for 3 h. UBR7/UBR box fused with N-terminal MYC and ATG8a/ATG8a^R13A^/ATG8a^3A^ fused with C-terminal HA were used and therefore detected by immunoblotting with the anti-MYC and anti-HA antibodies, respectively. Ponceau S (PS) and Coomassie brilliant blue (CBB) staining served as loading controls. (G) BiFC assay for *in vivo* interaction between ATG8a and UBR box. GFP^N^, GFP^C^, and their fusions with UBR box, ATG8a, ATG8a^R13A^, and ATG8a^3A^ were co-expressed in Col-0 protoplasts, followed by treatment with MG132 (10 μM) for 3 h. Reconstituted GFP fluorescence was visualized under a confocal microscope. DAPI staining indicates the location of nuclei. UBR, UBR box; Chl, chlorophyll; BF, bright field. Bars, 10 μm.

**Figure S3.**
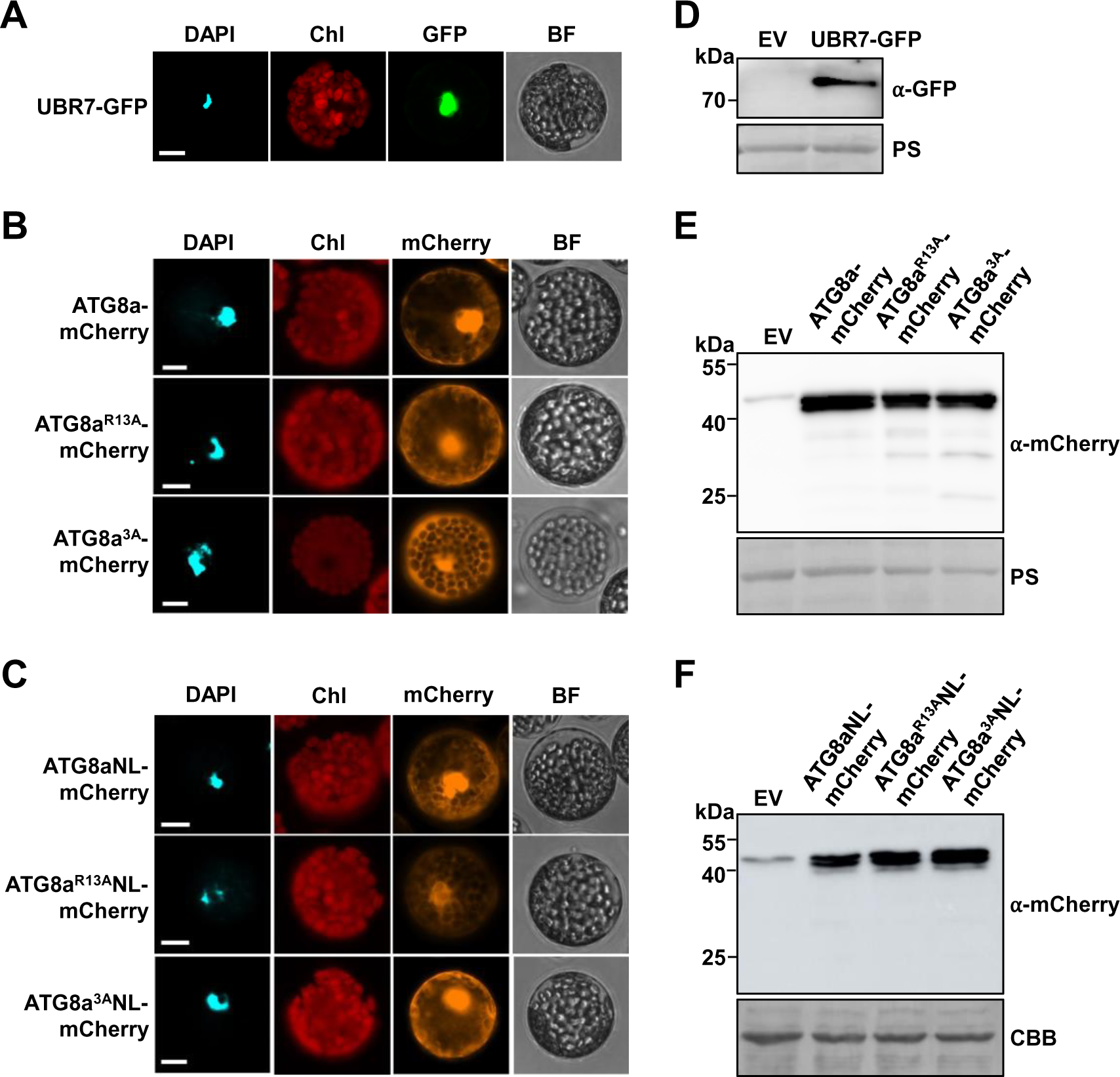
UBR7 and ATG8a are co-localized in the nucleus, related to Figure 3. (A-C) Subcellular localization of UBR7 (A), ATG8a (B), and ATG8aNL (C). UBR7-GFP, ATG8a/ATG8a^R13A^/ATG8a^3A^-mCherry, and ATG8aNL/ATG8a^R13A^NL/ATG8a^3A^NL-mCherry were expressed in Col-0 protoplasts as indicated. Fluorescence signal was visualized under a confocal microscope. DAPI staining indicates the location of nuclei. Chl, chlorophyll; BF, bright field. Bars, 10 μm. (D-F) Expression of UBR7 (D), ATG8a (E), and ATG8aNL (F) in transfected protoplasts. Protein lysates were prepared from Col-0 protoplasts transfected with the indicated constructs, followed by treatment with MG132 (10 μM) for 3 h. UBR7 and ATG8a proteins were detected by immunoblotting with the anti-GFP and anti-mCherry antibodies, respectively. Ponceau S (PS) and Coomassie brilliant blue (CBB) staining served as loading controls. EV, empty vector.

**Figure S4.**
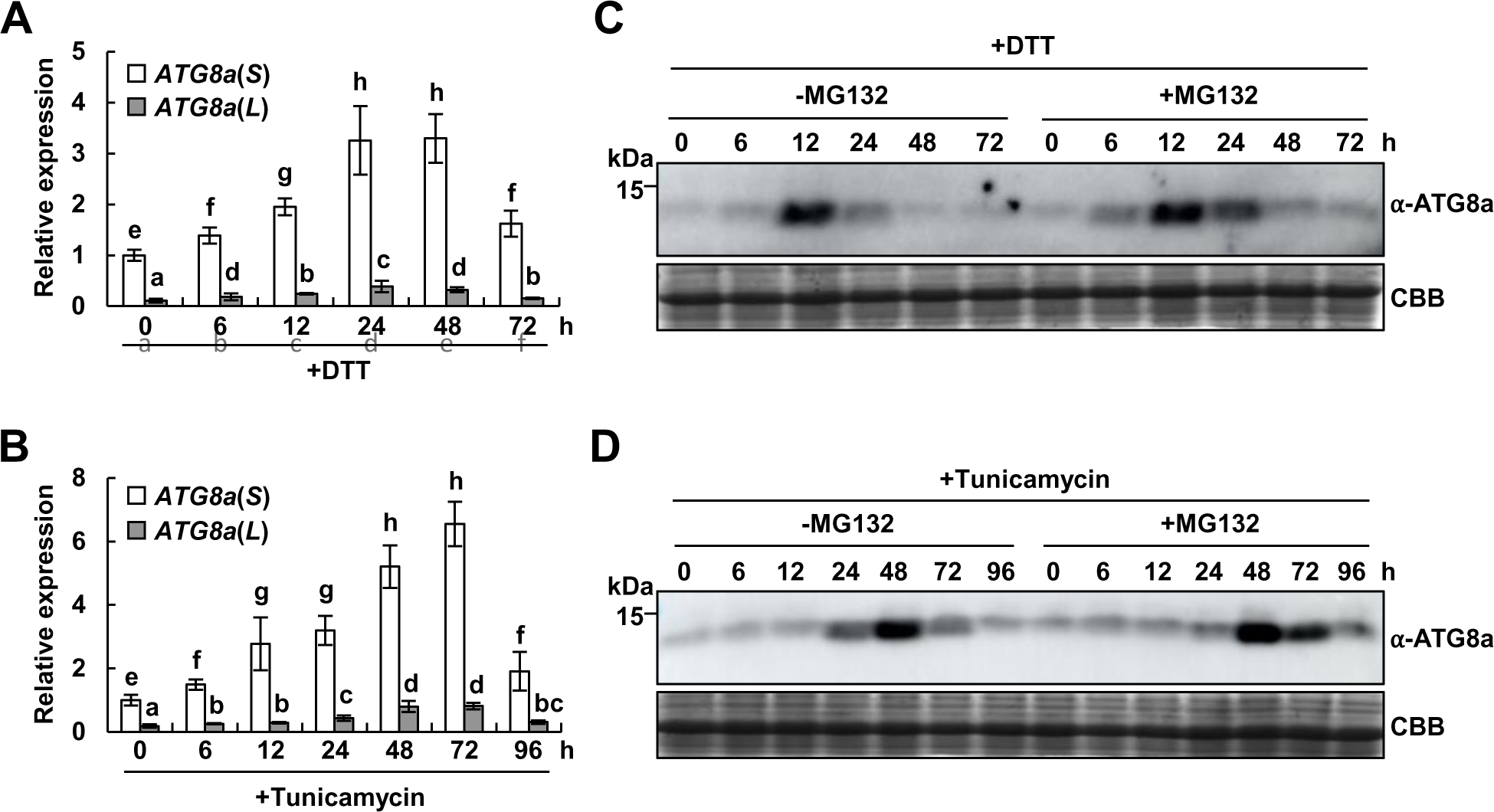
N-degron-mediated ATG8a degradation is not related to ER stress responses, related to Figure 4. (A and B) *ATG8*(*S*) expression is gradually increased in response to ER stressors, DTT (A) and tunicamycin (B). Transcript levels of *ATG8*(*S*) and *ATG8a*(*L*) were analyzed by RT-qPCR in Col-0 plants treated with DTT (8 mM) and tunicamycin (250 ng/ml) for the indicated times. Data represent means ± SD (*n* = 4 biological replicates). Different letters indicate significant differences (Tukey’s HSD test; *P* < 0.05). (C and D) ATG8a protein levels are not altered by MG132 treatment under ER stressors, DTT (C) and tunicamycin (D). ATG8a expression was monitored in the presence or absence of MG132 (10 μM) in Col-0 plants treated with DTT (8 mM) and tunicamycin (250 ng/ml) for the indicated times. Total proteins were extracted from plants and analyzed by immunoblotting with the anti-ATG8a antibody. Coomassie brilliant blue (CBB) staining served as a loading control.

**Figure S5.**
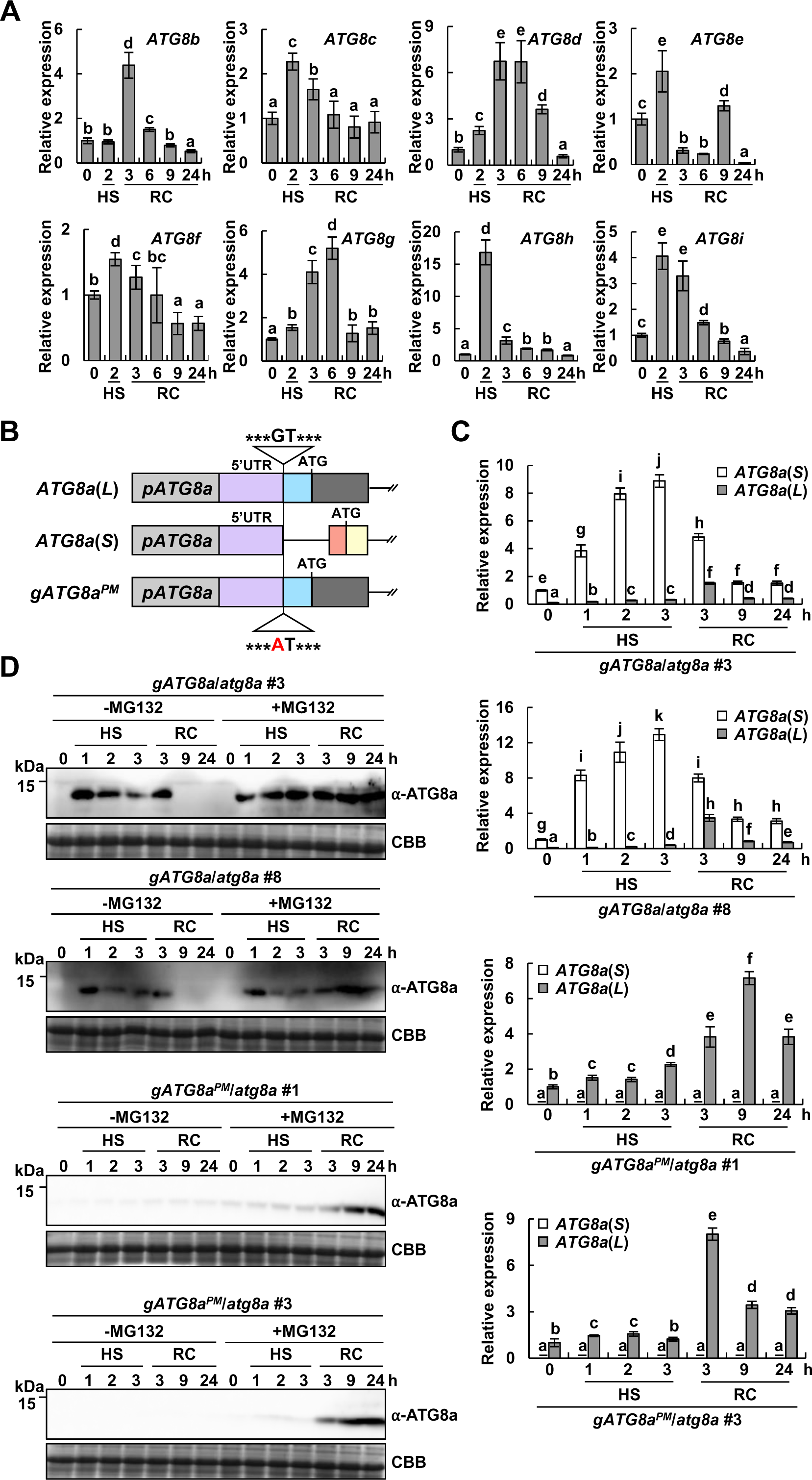
Expression analysis of ATG8 isoforms in wild-type and *atg8a* complemented with *ATG8a* genomic DNAs with either the wild-type sequence (*gATG8a*/*atg8a*) or a splice site mutation (*gATG8a^PM^*/*atg8a*), related to Figures 5 and 6. (A) Expression of several *ATG8* genes is induced during the HS recovery phase. Transcript levels of *ATG8* genes were analyzed by RT-qPCR in Col-0 plants exposed to HS (42°C) for 2 h and recovery (23°C) for the indicated times. Data represent means ± SD (*n* = 4 biological replicates). Different letters indicate significant differences (Tukey’s HSD test; *P* < 0.05). (B) Genomic structures of *ATG8*(*L*), *ATG8a*(*S*), and *gATG8a^PM^* with a G to A splice site mutation in the retained intron. Exons are indicated by yellow and dark gray (coding) and purple, red, and blue (UTRs) boxes. Introns are shown in lines. Triangles indicate the position of a G to A splice site mutation. (C) *ATG8*(*S*) is barely expressed in *gATG8a^PM^*/*atg8a* lines. *ATG8*(*S*) and *ATG8a*(*L*) expression were analyzed by RT-qPCR in *gATG8a*/*atg8a* and *gATG8a^PM^*/*atg8a* lines exposed to HS (42°C) and recovery (23°C) for the indicated times. Data represent means ± SD (*n* = 4 biological replicates). Different letters indicate significant differences (Tukey’s HSD test; *P* < 0.05). (D) ATG8a expression is decreased in *gATG8a^PM^*/*atg8a* lines under HS. ATG8a expression was monitored in the presence or absence of MG132 during HS and recovery in *gATG8a*/*atg8a* and *gATG8a^PM^*/*atg8a* lines. For MG132 treatment, plants were sprayed with MG132 (10 μM) prior to exposure to HS and recovery conditions. Total proteins were extracted from plants exposed to HS (42°C) and recovery (23°C) for the indicated times and analyzed by immunoblotting with the anti-ATG8a antibody. Coomassie brilliant blue (CBB) staining served as loading controls.

**Figure S6.**
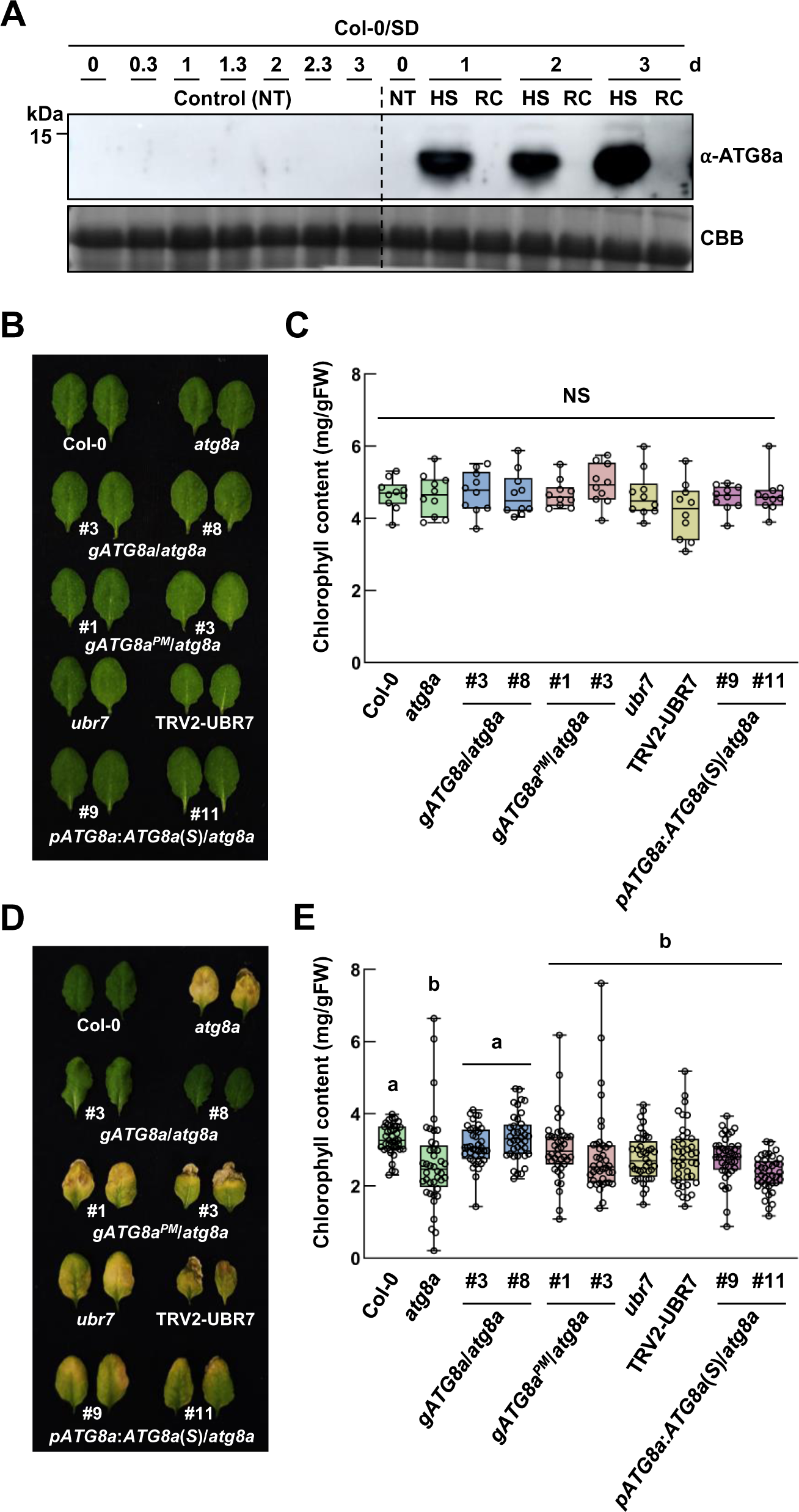
ATG8a turnover is important for plant thermotolerance, related to Figure 6. (A) Fluctuations in ATG8a abundance coincide with daily HS and recovery cycles. Col-0 plants grown in short days were exposed to 42°C (HS) for 8 h during the day and maintained at 23°C (recovery) for the remaining 16 h. Total proteins were extracted from Col-0 plants exposed to HS and recovery repeatedly for 3 days and analyzed by immunoblotting with the anti-ATG8a antibody. Coomassie brilliant blue (CBB) staining served as a loading control. SD, short day; NT, no treatment; HS, heat stress; RC, recovery. (B) Leaf phenotypes of Col-0, *atg8a*, *ubr7*, TRV2-UBR7, and complementation lines grown in short days. Plants were maintained at 23°C under short-day conditions during HS experiments. (C) Quantification of chlorophyll content in control plants in (B). Data represent means ± SD (*n* = 10 leaves). NS, not significant. (D) Thermotolerance phenotypes of Col-0, *atg8a*, *ubr7*, TRV2-UBR7, and complementation lines under recurring HS and recovery conditions. Plants grown in short days were exposed to 42°C (HS) for 8 h during the day and maintained at 23°C (recovery) for the remaining 16 h, which was repeated for 5 days. Plants were allowed to recover for an additional 3 days and observed for leaf phenotypes. (E) Quantification of chlorophyll content in plants exposed to HS and recovery in (D). Data represent means ± SD (*n* = 40 leaves). Different letters indicate significant differences (Tukey’s HSD test; *P* < 0.05).

**Figure S7.**
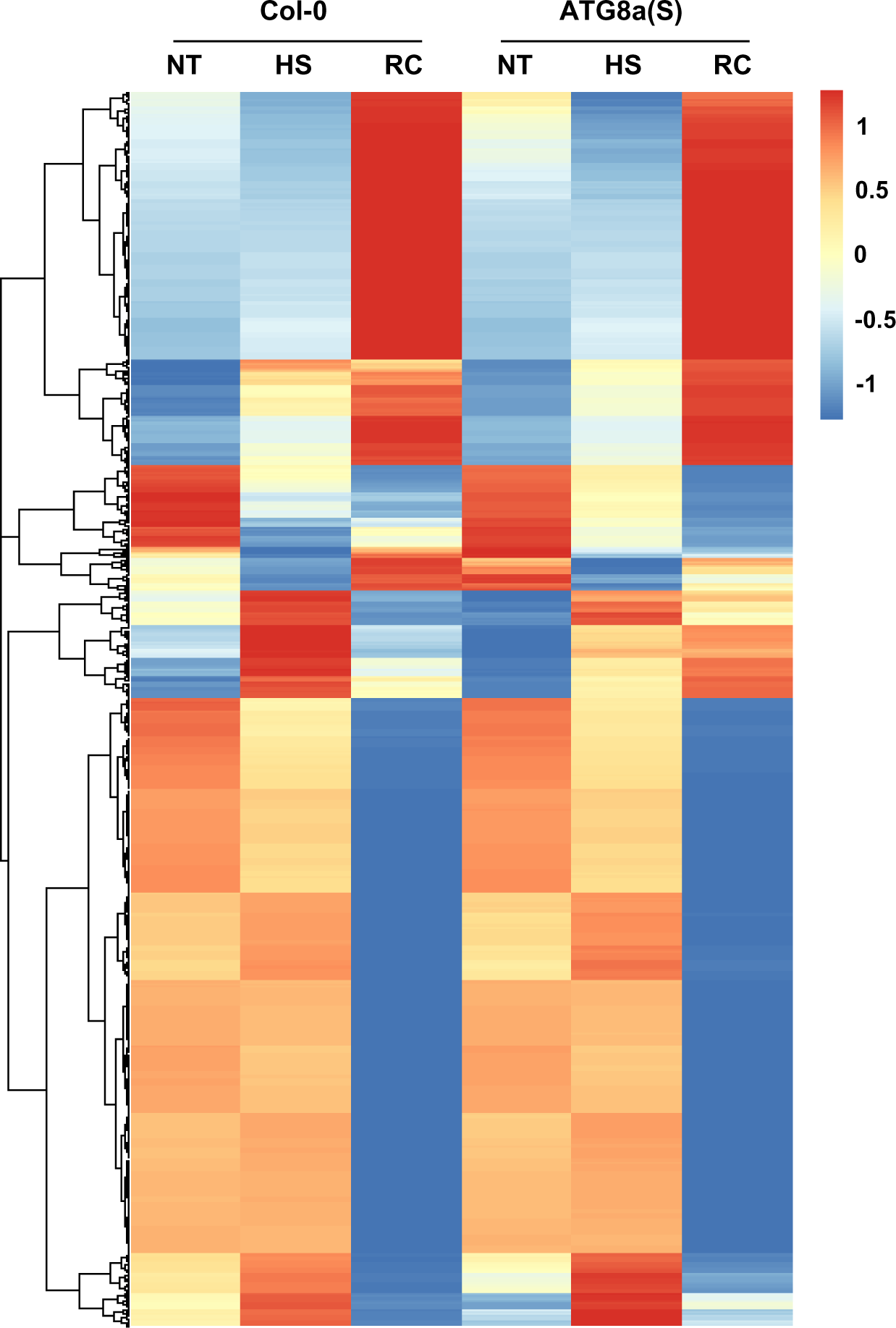
Heatmap of proteomics profiles in Col-0 and *pATG8:ATG8a*(*S*)/*atg8a* plants under NT, HS, and RC conditions, related to Figure 7. Heatmap was generated using the pheatmap package in R Studio. NT, no treatment; HS, heat stress; RC, recovery.

## Notes

### Competing Interest Statement

The authors have declared no competing interest.

